# Natural microbial exposure imposes layered constraints on epithelial and type 2 immunity

**DOI:** 10.64898/2026.05.04.722698

**Authors:** D. Ovezgeldiyev, R. Doolan, V. Trefzer, A. Baltensperger, S. Rezaei, A. Serra, C. Pohl, N. Putananickal, B. Chalethu, E. Kuku, J. Dommann, P. Schneeberger, S Runge, V Lang, C Gmeiner, C Günther, S. P. Rosshart, T. Bouchery

## Abstract

The tuft cell-ILC2 amplification circuit has emerged as a central paradigm of anti-helminth immunity in laboratory animals, driving the so-called weep and sweep response, required for parasite expulsion. Yet soil-transmitted helminths (STHs) commonly establish chronic infections in humans. Whether tuft cell expansion is required for parasite clearance under naturalistic conditions remains unknown. Here, using wildlings, a naturalized mouse model exposed from birth to complex microbial communities and pathogens, we re-investigated anti-STH immunity in the context of ecological realism. Following infection with *Nippostrongylus brasiliensis*, specific pathogen-free (SPF) mice mounted markedly amplified type 2 responses in both lung and intestine compared to wildlings. In the intestine, SPF mice mounted robust tuft cell expansion, IL-25 production, ILC2 accumulation, and goblet cell hyperplasia. In contrast, infected wildlings exhibited delayed parasite expulsion, limited tuft cell expansion, reduced IL-25 and ILC2 responses, and attenuated goblet cell expansion. Wildling tuft cells, but not goblet cells, displayed markedly reduced expansion in response to succinate or exogenous IL-13, indicating selective hypo-responsiveness of the epithelial sensory compartment. Microbial transfer into adult SPF mice selectively conferred tuft cell hypo-responsiveness without impairing ILC2 accumulation or goblet cell expansion. Tuft cell hypo-responsiveness in wildlings and FMT recipients was associated with enrichment of fermentative bacteria and increased levels of the short chain fatty acids acetate and propionate. Together, these findings indicate that ecological microbial exposure imprints systemic type 2 immunity during early-life, whereas epithelial responsiveness remains plastic and microbiome-dependent, thereby revealing regulatory constraints not evident under SPF conditions.

**One Sentence Summary:** Our findings reveal that in a naturalized immune-microbiome context, intestinal tuft cells are surprisingly hypo-responsive, highlighting how environmental microbial exposure can calibrate type 2 immunity and helminth resistance.

## INTRODUCTION

Laboratory mice have provided vital insight into mammalian immune systems, yet their immune responses often differ strikingly from those of wild mammals^1^. Whether these differences reflect exaggerated responses in pathogen-limited environments remains unclear.

Soil-transmitted helminths (STHs) contribute to perpetuating a cycle of poverty in vulnerable populations^2^. The major human STHs - *Ascaris lumbricoides, Trichuris trichiura*, and hookworms (principally *Necator americanus*) - primarily infect individuals living in impoverished settings with limited access to sanitation and clean water. These infections are clinically associated with malnutrition, anemia, stunted growth, and impaired cognitive development in children, thereby exerting a profound impact on human capital. Globally, STH infections account for an estimated 3.5 to 4.9 million Disability-Adjusted Life Years (DALYs) lost annually^3^, a burden comparable to or exceeding that of other tropical diseases that receive greater public health attention.

Human STH infections induce pronounced eosinophilia, elevated IgE, and a mixed cytokine profile including Th1, regulatory, and anti-helminth Th2 responses. Murine models have demonstrated that type 2 immune responses are protective, both in primary infections, via the “weep and sweep” expulsion of STH, and in secondary infections, where M2 macrophages contribute to parasite trapping and killing^4–10^. Yet, humans experience chronic infection, rapid reinfection and generally non-sterilizing immunity after natural STH exposure^11,12^. This epidemiological reality stands in stark contrast to the frequently protective outcomes observed in conventional laboratory models and complicates identification of immune correlates that will translate to the field. Here, we ask which immune mechanisms remain protective under natural microbial pressure.

As a result of their long co-evolution, bi-directional interactions between microbiota and helminths further complicate this picture. For example, eggs of *Trichuris* spp. require bacterial cues to initiate development, and helminths can reshape the microbiome to create a favorable niche^13–16^. Given this close proximity, we reasoned that SPF mice, lacking natural microbial and pathogen exposure, might not reflect accurately the niche in which STH normally develop in wild mammals and might exaggerate type 2 responses, obscuring physiologically relevant immune mechanisms.

“Naturalization” strategies have been developed to bridge this gap, including adult colonization of laboratory mice with wild microbiota or housing in outdoor enclosures and are starting to be applied to STHs^1,17^. For example, “rewilded” mice in an outdoor enclosure acquired a more Th1 dominant immune profile and became susceptible to *T. muris* infection^18^. These approaches, however, typically miss early-life exposure and lifelong pathogen encounters, which are increasingly recognized as a critical window for immune education.

To explore how complex microbial and pathogen exposure affects parasite establishment and host immunity, we choose to use wildling mice^19–21^. Those mice are exposed to a diverse community of bacteria, viruses, protozoa, fungi, and mites *in utero* and throughout life. This approach is stable over years, reproducible, and allows us to house wildling and SPF counterparts side-by-side under identical conditions, minimizing environmental confounders^1,21^. Here, we utilized *N. brasiliensis* infection, a parasite which is widely used to investigate type-2 immunity and has been centrally involved in defining the mucosal response, allowing us to ask how early-life microbial and pathogen exposure recalibrates type-2 immunity during helminth infection.

## RESULTS

### Subhead 1: Microbial experience preserves parasite establishment but limits lung tissue injury

To determine whether ecological microbial exposure alters early events of helminth infection, we infected male and female wildlings and SPF counterparts with 500 L3 *N. brasiliensis* larvae subcutaneously **(Fig 1A)**. Establishment of the parasite in the lungs (2 days post-infection, 2 dpi) and subsequent migration to the intestine (5 days post infection) were comparable between microbiome-experienced and SPF mice in both sexes **(Fig. 1B&C and S1A&B),** indicating that microbial conditioning does not impair infectivity or early parasite development. Consistent with this, larval molting and size in the lungs were similar between groups **(Fig. S1C)**, excluding developmental differences.

**Fig. 1.**
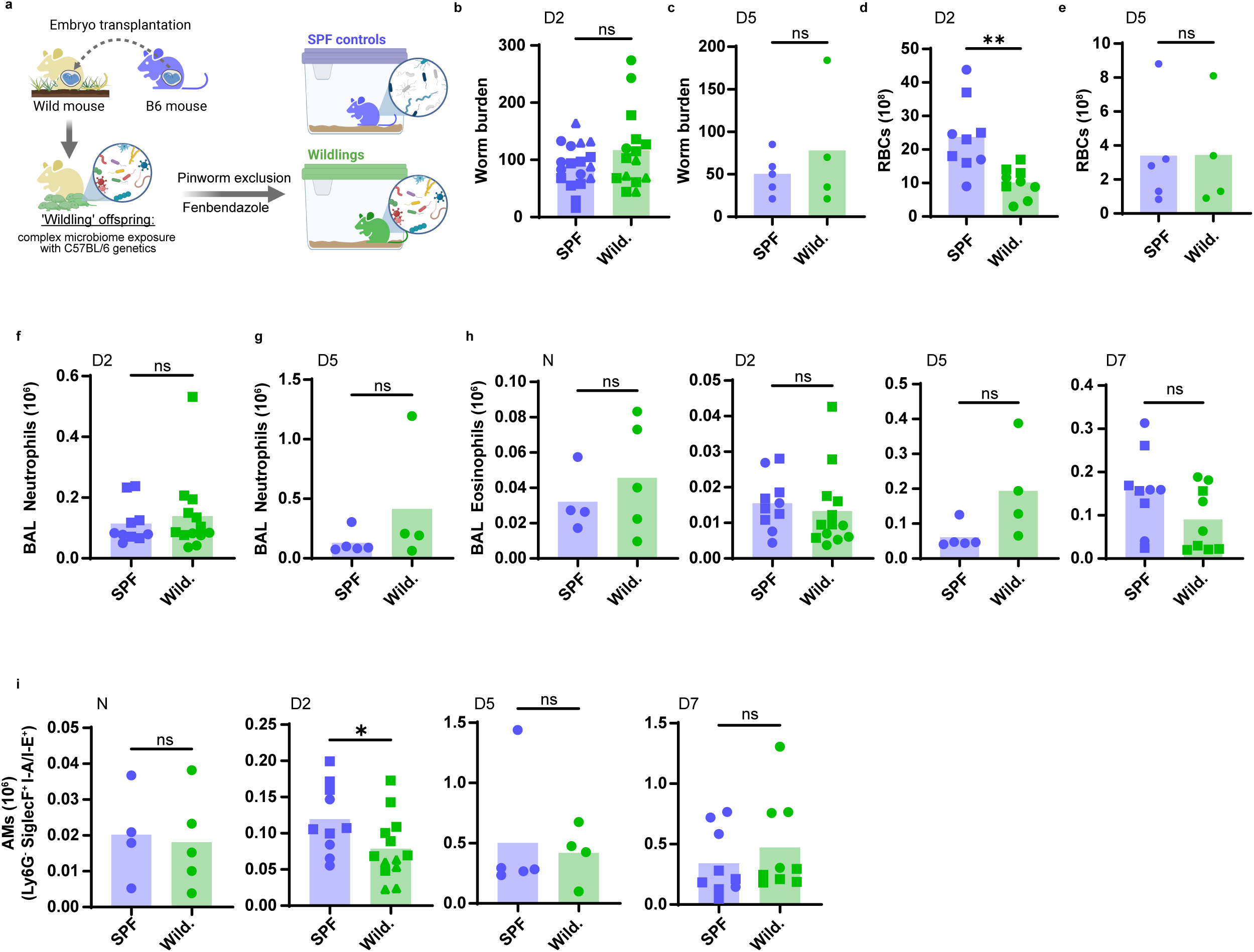
*N. brasiliensis* establishment in the lungs of wildlings mirror SPF response. Wildlings and SPF mice as depicted in **(A)** were infected by subcutaneous infection with 500 L3 of *N. brasiliensis,* for 0, 2, 5 or 7 days (Naïve, D2-7). Worm burden was measured by baermann of the (**B**) lungs 2 days post-infection and in the (**C**) small intestine 5 days post-infection. At both times, bronchoalveolar lavage of the lungs was performed, and red blood cells (RBCs) counted to assess hemorrhaging (**D**) and (**E**). Flow cytometry of BAL samples was performed to quantify the total number of immune cell subpopulations in the airways of each mouse. CD45^+^ Ly6G^+^ Gr1^+^ neutrophils were quantified at D2 and D5 post-infection (**F**) and (**G**) respectively. CD45^+^ LyG^−^ SiglecF^+^ I-A/I-E^+^ alveolar macrophages (AMs) (**I**) and CD45^+^ LyG^−^ SiglecF^+^ I-A/I-E^−^ SCC^high^ eosinophils **(J)** were quantified in naive uninfected (N) mice, and at 2, 5 and 7 days post-infection (D2-7). Each dot represents one mouse. D2 and D7 time points are pooled from 3 independent experiments (n=4-5); D5 and Naive are issued from 1 experiment (n=4-5). When possible, different symbols are used between biological repeats.

During its migration through the lungs, *N. brasiliensis* causes tissue damage and hemorrhage. Interestingly, despite equivalent worm burden, lung hemorrhage at 2 dpi was markedly reduced in wildlings compared to SPF mice **(Fig. 1D, S1D)**. In both types of mice, the hemorrhage was fully resolved by 5 days post-infection, as previously reported for SPF mice **(Fig. 1E)**.

Because hemorrhage following *N. brasiliensis* infection is typically associated with neutrophil infiltration and subsequent tissue damage, we quantified neutrophils in bronchoalveolar lavage fluid. Although neutrophil frequency was modestly increased in wildlings **(Fig. S1E)**, absolute cell numbers were comparable between microbiome conditions **(Fig. 1F)**, arguing against reduced neutrophil recruitment as the explanation for diminished hemorrhage. Neutrophil numbers declined similarly in both groups by 5 dpi **(Fig. 1G, S1F)**.

We next assessed additional hallmarks of lung inflammation. Eosinophil numbers in bronchoalveolar lavage were similar between SPF and wildlings, both at baseline and after infection **(Fig.1H, S1G),** indicating preserved early type 2 recruitment. In contrast, alveolar macrophages were transiently reduced in wildlings at 2 dpi, both in absolute number and as a proportion of live cells **(Fig. 1I, S1H),** suggesting a modest delay in macrophage expansion at the peak of tissue damage. By later time-points, absolute macrophage numbers were comparable between groups, although proportional differences persisted (**Fig. S1H**).

Altogether, we show that ecological microbial conditioning in mice preserves the infection establishment kinetic with a moderate attenuation of the early pulmonary inflammatory response and tissue damage.

### Subhead 2: Microbial experience restrains pulmonary type 2 amplification and skews immune programming

We next examined whether microbial experience alters the development of pulmonary type 2 immunity following *N. brasiliensis* infection. Because Th2 cells and ILC2s sustain alternatively activated macrophages (AAMs), we quantified these populations at day 10 post-infection, after parasite egress from the lungs. In naïve mice, Th2 cells and ILC2s were present at comparable frequencies in SPF and wildlings **(Fig. 2A&B)**, indicating similar basal type 2 tone. Following infection, both populations expanded robustly in SPF mice, whereas this expansion was markedly blunted in wildlings. Consequently, at day 10 post-infection, both Th2 and ILC2 numbers were significantly reduced in microbiome-experienced mice relative to SPF controls **(Fig. 2A&B)**, demonstrating impaired amplification of the type 2 compartment rather than a baseline deficiency.

**Fig. 2.**
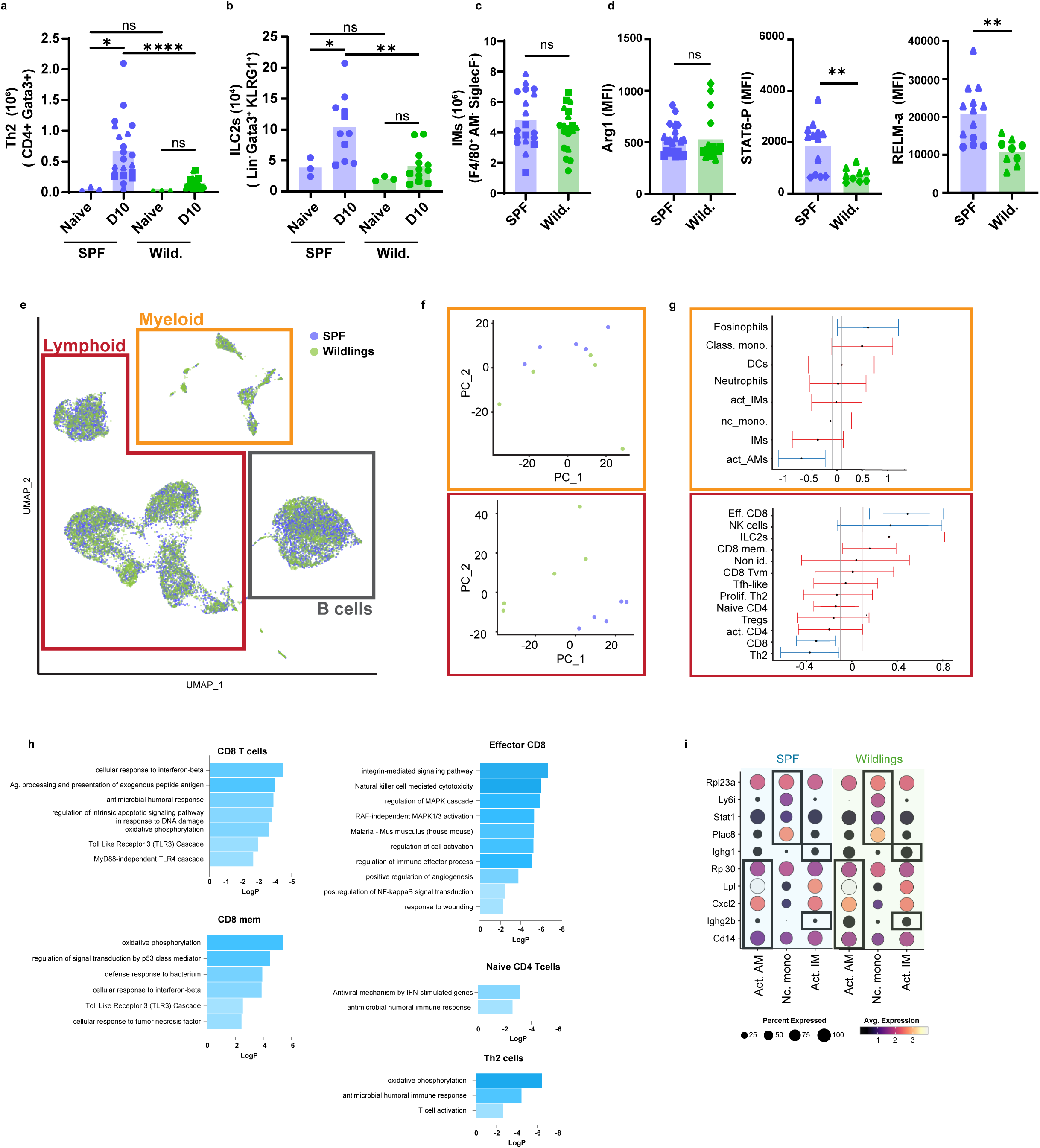
Wildlings have a dampened type-2 response and an increase in type-I IFN signature in the lungs. Wildlings and SPF mice were infected by subcutaneous infection with 500 L3 of *N. brasiliensis,* 10 days. Lungs cells were isolated by digestion from naïve or 10 days infected mice and analysed by flow cytometry **(A)** Number of Th2 cells (CD45^+^ CD4^+^ Gata-3^+^) **(B)** Number of ILC2s (CD45^+^ Lin^−^ KLRG1^+^ Gata-3^+^) Pool of 3 independent experiments, representative of one more (n=4-8) **(C)** Number of interstitial macrophages (CD45^+^ F4/80^+^ AM^−^ SiglecF^−^) **(D)** Mean Fluorescence Intensity of STAT6-P, RELMa and Arg-1 expression gated on interstitial macrophages. Pool of 2 independent experiments, representative of 2 more independent experiments (n=4-8). **(E)** Single cell RNA sequencing was conducted on cell isolated from infected lungs 10 days post-infection (n=4, per mouse type). UMAP visualization of all cells clustered by Seurat in wildlings and SPF mice. Cells were sorted in three subsets “B cells”, “myeloid” and “lymphoid” for further analysis. **(F)** PCA of cells in the “myeloid” and “lymphoid” subsets **(G)** Bar plot showing the percentages of cells for each subset. Blue indicates a significant change between Wildling and SPF condition. **(H)** Over-representation analysis showing enriched pathways of the cells in the “lymphoid” subset for comparison of wildlings versus SPF conditions, performed in Metascape (DEGs selected at p=0.05). **(I)** Bubble plot of expression level of differentially expressed genes for monocytes and macrophages for comparison of wildlings versus SPF conditions. When possible, different symbols are used between biological repeats.

Given the established dependence of interstitial macrophage polarization on type 2 cytokines, we next assessed macrophage activation status. The number of interstitial macrophages was not affected by the microbial experience of the mice **(Fig. 2C)**, indicating proliferation and recruitment are likely not affected. However, markers of alternative activation were selectively diminished in wildlings: phosphorylation of STAT6 and expression of RELM-α were significantly reduced **(Fig. 2D)**, consistent with attenuated M2 polarization. Thus, restrained type 2 lymphocyte expansion in wildlings translates into functional consequences for macrophage programming.

To obtain an unbiased view of these alterations, we performed single-cell RNA sequencing (BD Rhapsody) on total lung cells at day 10 post-infection (8,633 SPF and 9,536 wildling cells across five mice per group). Global UMAP visualization revealed broadly similar cellular architecture between groups **(Fig. 2E, Fig. S2A, Table 1)**, suggesting that microbial experience does not grossly remodel lung composition at this stage. As previously reported, the B cell compartment showed the most pronounced transcriptional divergence^21^. Pathway enrichment analysis highlighted signatures associated with “B cell receptor signaling,” “defense response to bacterium,” and “response to lipopolysaccharide” in wildlings **(Fig. S2A)**, consistent with heightened microbial exposure.

To resolve whether differences were concentrated within specific immune compartments, we analyzed lymphoid and myeloid subsets separately **(Fig. S2B&C)**. Principal component analysis indicated that inter-group variability was driven predominantly by the lymphoid compartment **(Fig. 2F)**. Consistent with flow cytometry data, Th2 cells were proportionally reduced in wildlings **(Fig. 2G)**. Due to limited cell numbers, ILC2 representation could not be robustly assessed by scRNA-seq. In contrast, wildlings displayed increased proportions of effector CD8 T cells and NK cells, in line with prior characterization of this model. Pathway enrichment analysis in CD8 T cells and naïve CD4 T cells revealed enhanced interferon and antibacterial response signatures in wildlings **(Fig. 2H**, **Table 2)**. Notably, Th2 cells from wildlings showed enrichment in pathways related to “antimicrobial humoral immune response” and “oxidative phosphorylation,” suggesting altered functional and metabolic programming rather than simple expansion reduction.

In the myeloid compartment, only 28 differentially expressed genes were identified across all clusters between groups, indicating comparatively modest transcriptional divergence. Enrichment analysis in monocyte and macrophage subsets revealed limited pathway differences, but again with signatures consistent with antibacterial responsiveness in wildlings, including “defense response to bacterium,” “mononuclear cell proliferation,” and “response to lipopolysaccharide” **(Fig. 2I)**.

Collectively, these data indicate that microbial experience does not abrogate type 2 *immunity per se* but constrains its amplification and reprograms the pulmonary immune environment toward an interferon- and antibacterial-biased state. This shift dampens alternative macrophage polarization and suggests that prior microbial conditioning establishes a competing inflammatory axis that modulates helminth-induced type 2 responses.

### Subhead 3: Microbial experience delays helminth expulsion by restraining the epithelial-type 2 circuit

Having observed restrained type 2 amplification in the lung of microbiome-experienced mice, we next asked whether this reflected a tissue-specific phenomenon or a broader recalibration of barrier immunity. We therefore examined intestinal clearance of *N. brasiliensis*, which depends on coordinated epithelial and type-2 immune activation, via the canonical “weep and sweep” response. At day 7 post-infection, corresponding to the early expulsion phase, wildlings retained significantly higher adult worm burdens than SPF mice **(Fig. 3A, left)**. By day 10, all SPF mice had cleared the infection, whereas a subset of wildlings continued to harbor substantial worm loads **(Fig. 3A, right)**, accompanied by increased egg counts in the cecum **(Fig. 3B)**. Nevertheless, by day 20 post-infection, parasites were undetectable in both groups, indicating that microbial experience delays but does not prevent self-cure. This delayed expulsion phenotype was observed in both males and females **(Fig. S3A)**.

**Fig. 3.**
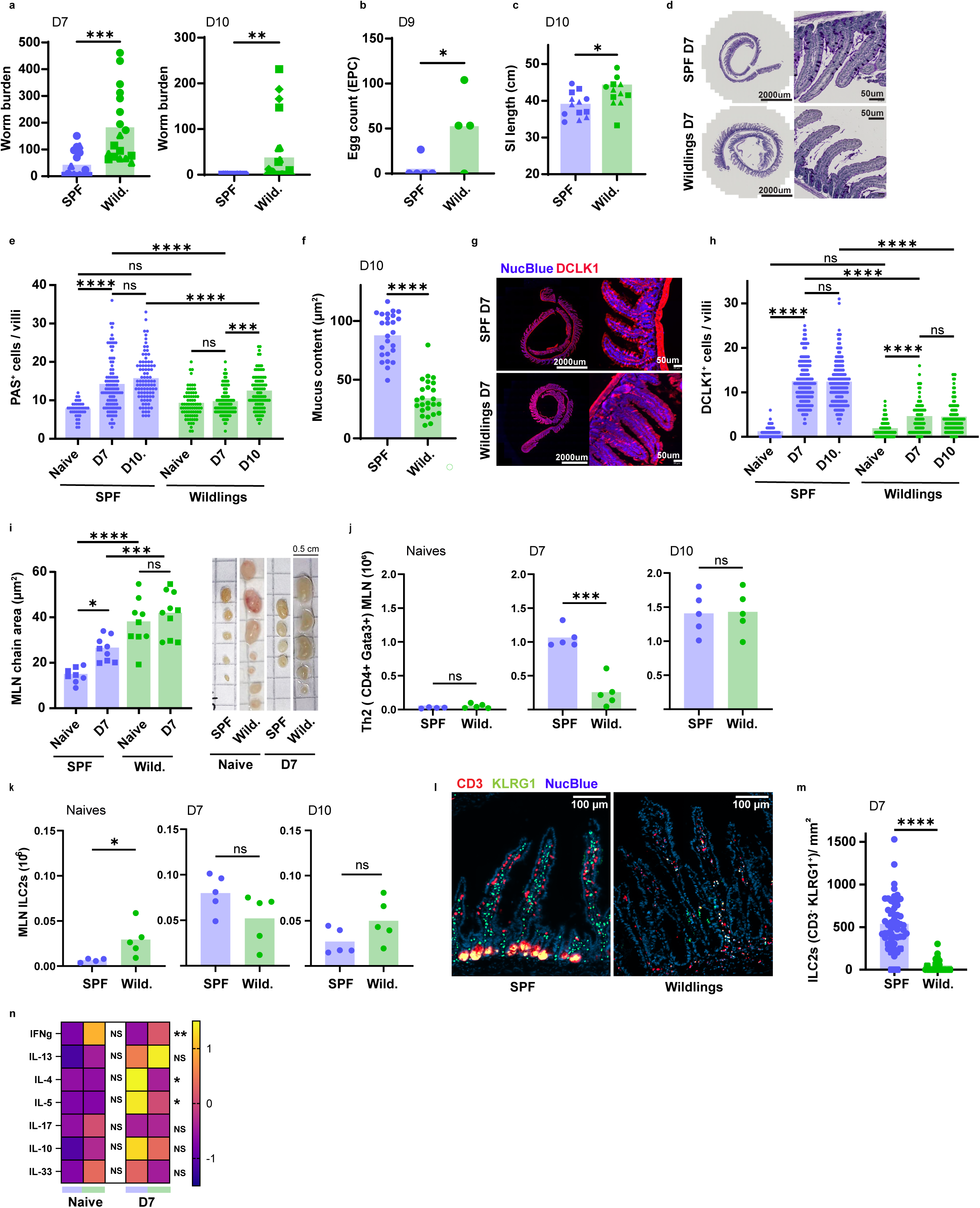
Wildlings have a delayed intestinal expulsion of *N. brasiliensis*. Wildlings and SPF mice were infected by subcutaneous infection with 500 L3 of *N. brasiliensis, 7-*10 days. Worm burden was measured by baermann of the (**A**) small intestine 7-9 (early) or 10-11 (late) days post-infection. Data pooled from 3 independent experiments (n= 4-8) **(B)** Eggs were counted in the caecum content and expressed by total volume of content. Data representative of 2 independent experiments (n=4) **(C)** Small intestinal length was measured during late expulsion (D10). 3 pooled experiment (n=4-8) **(D-F)** PAS staining of Swiss-roll of 3 anterior cm of small intestine. Representative images showing goblet cell hyperplasia in SPF mice (D). (E) Quantification of the number of Goblet cells per villi on sections described in (D) and on the area of PAS+ cells (F). Pool of 2 independent experiments per time point (n=4-6) **(G&H)** Representative images (G) and tuft cell quantification (H) by immunofluorescence analysis of DCLK1 in the small intestine from naïve or infected mice. **(I)** Representative images and Mesenteric Lymph Node (MLN) chain size quantification between wildlings and SPF, in naïve mice or mice 9 days post-infection. Pool of 2 independent experiments (n=4-6). **(K&L)** Number of Th2 cells (K, CD45^+^ CD4^+^ Gata-3^+^) and ILC2s (L, CD45^+^ Lin^−^ KLRG1^+^ Gata-3^+^) in the MLN of naïve mice or mice in early or late phase of expulsion. 1 experiment, representative of 2 more (but on partial chain of MLN; n=4-5). **(M)** Quantification by immunohistofluorescence of the number of ILC2 per surface of small intestinal villi (CD3-KLRG1+) in SPF or wildling mice infected with Nb for 7 days. **(N)** Intestinal cytokines measured by MSD in naïve or infected mice in SPF or wildlings for 7 days. 1 experiment (n=3-6 mice). When possible, different symbols are used between biological repeats.

Because helminth clearance is driven primarily by epithelial effector mechanisms, we next examined intestinal physiology. wildlings were leaner and exhibited a significantly longer small intestine post-infection and a slight increase in villi length **(Fig. S3B&C, Fig. 3C)**, suggesting intrinsic differences in intestinal architecture.

Baseline goblet cell numbers were comparable between groups **(Fig. 3D&E)**. Following infection, SPF mice showed the expected robust goblet cell hyperplasia by day 7, which plateaued thereafter. In contrast, wildlings failed to significantly increase goblet cell numbers at the early expulsion time point; hyperplasia became evident only later, consistent with delayed worm clearance. At all post-infection timepoints, goblet cell numbers remained lower in wildlings compared to SPF controls. Moreover, goblet cell mucus content was reduced in wildlings during the expulsion phase **(Fig. 3F)**, indicating constrained epithelial proliferation and activation.

Upstream of goblet cells, tuft cells act as sensory sentinels that initiate type 2 amplification via IL-25 release^6^. At baseline, wildlings exhibited a modest increase in DCLK1 tuft cells compared to SPF mice **(Fig. 3G&H)**. Upon infection, SPF mice mounted a strong tuft cell expansion by day 7, whereas wildlings displayed only a moderate increase that plateaued at significantly lower levels.

To test whether downstream type 2 immunity was affected, we first assessed mesenteric lymph nodes (MLNs) as a surrogate for intestinal responses. Wildlings displayed increased MLN number and total chain area at baseline compared to SPF controls, and this did not further expand upon infection **(Fig. 3I)**. In contrast, SPF mice exhibited the expected MLN enlargement following infection. Flow cytometry revealed a delayed and dampened Th2 expansion in wildlings **(Fig. 3J)**, whereas total ILC2 numbers in MLNs were comparable between groups **(Fig. 3K)**.

Because MLN analysis might underestimate tissue-resident responses, we next quantified ILC2s directly in the lamina propria at day 5 post-infection, coinciding with initiation of epithelial effector responses in SPF mice. KLRG1 ILC2s were significantly reduced in the lamina propria of wildlings relative to SPF controls **(Fig. 3L&M)**. Surprisingly, total IL-13 level in the intestinal tissue was not different between SPF and wildlings at baseline or 7 dpi **(Fig. 3N),** while both IL-4 and IL-5 levels were decreased in wildlings after infection **(Fig. 3N).**

Taken together, these findings indicate that microbial experience delays helminth expulsion by constraining the epithelial-type-2 amplification loop. Reduced tuft cell expansion and IL-25 production restrain local ILC2 accumulation, leading to impaired goblet cell hyperplasia and delayed execution of the “weep and sweep” program. Despite reduced ILC2 numbers and diminished IL-4 and IL-5 production, IL-13 levels were preserved in wildlings, suggesting that epithelial responses occur in the presence of sufficient type 2 cytokine signals. Clearance ultimately occurs, but with reduced magnitude and delayed kinetics, consistent with a recalibrated type 2 response in microbiome-experienced mice in both lungs and intestine.

### Subhead 4: Microbial experience selectively attenuates IL-13-driven tuft cell, but not goblet cell responses

Having observed attenuated tuft cell expansion during *N. brasiliensis* infection in wildlings, we hypothesized that microbial experience raises the activation threshold of the tuft-ILC2 amplification circuit. Tuft cells sense luminal cues through chemosensory pathways, including bitter taste receptors and metabolite sensors such as SUCNR1 and GPR41/43^22^. We therefore asked whether providing an exogenous metabolite stimulus could overcome the restrained epithelial response.

Mice received sodium succinate^23^ in drinking water from the day of infection until analysis **(Fig. 4A)**. As expected, succinate supplementation did not further reduce worm burden in SPF mice, which had already efficiently cleared infection by the early expulsion phase **(Fig. 4B)**. Strikingly, succinate failed to accelerate parasite clearance in wildlings, which suggests that tuft cells hypo-responsiveness is not specific to helminth infection. Consistent with prior reports, succinate increased tuft cell numbers per villus in SPF mice^23^. In wildlings, however, tuft cell expansion remained significantly blunted, even with supplementation **(Fig. 4C)**.

**Fig. 4.**
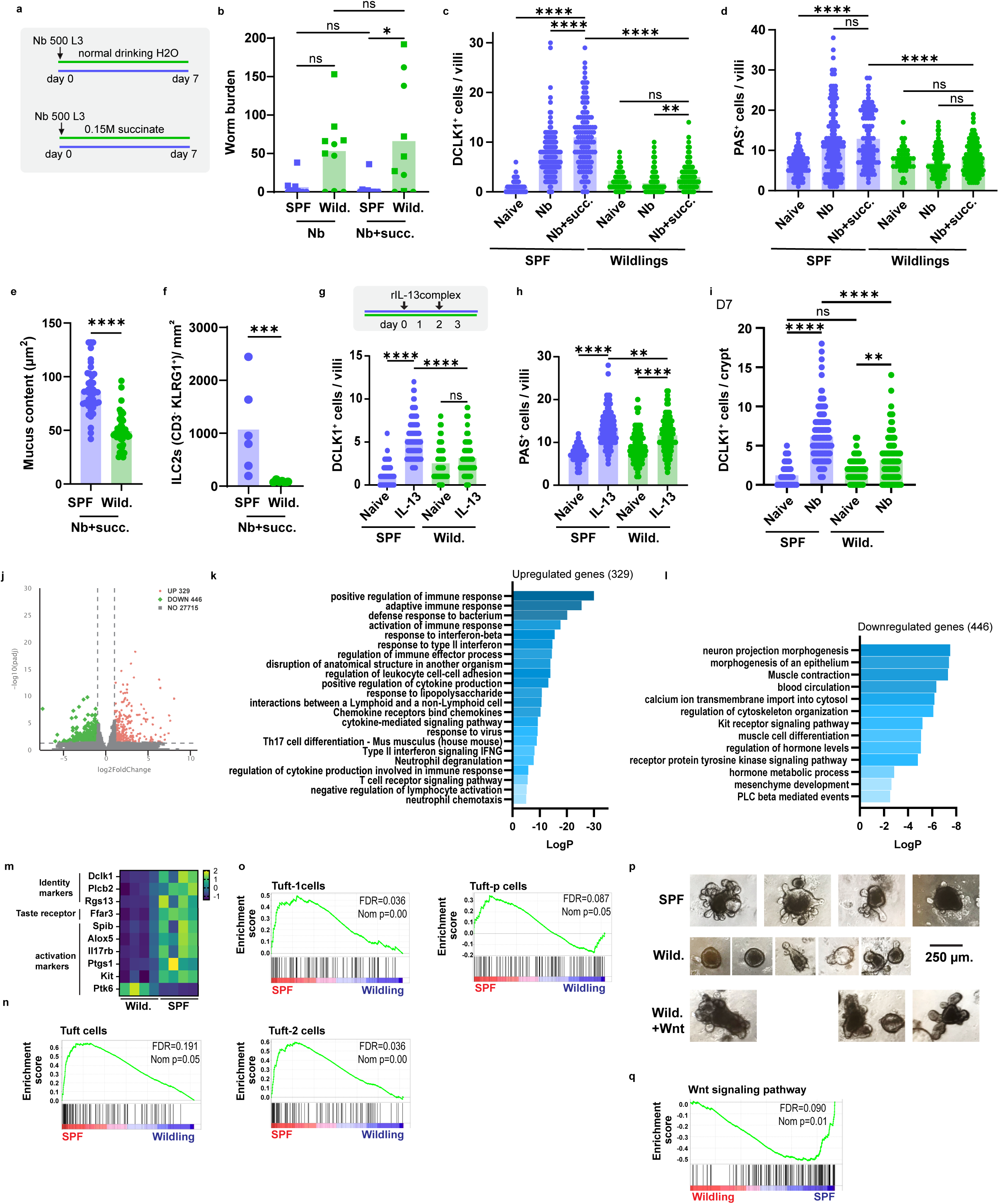
Tuft and goblet cells are differentially tuned to the same cytokine environment in wildlings. **(A-F)** Wildlings and SPF mice were infected by subcutaneous infection with 500 L3 of *N. brasiliensis, 7-*9 (early) days. Mice were further supplemented or not in the drinking water with succinate. **(B)** Worm burden was measured by baermann in the small intestine. Data pooled from 2 independent experiments (n= 5). **(C)** Tuft cell quantification by immunofluorescence analysis of DCLK1 in the small intestine from naïve or infected mice, per villi (C) **(D-E)** PAS staining of Swiss-roll of 3 anterior cm of small intestine. Quantification of the number of Goblet cells per villi (D) or of the area of Goblet cells (E). Data pooled from 2 independent experiments (n= 5). **(F)** Intestinal ILC2 number quantified by immunohistology (CD3-KLRG1+). **(G-H)** Naïve wildlings or SPF mice were treated with cIL-13 intraperitoneally or left untreated. Tuft cell quantification by immunofluorescence analysis of DCLK1. **(H)** Quantification of the number of Goblet cells per villi. 1 experiment (n=3-4), representative of 2 independent experiments. **(I)** Tuft cell quantification by immunofluorescence analysis of DCLK1 in the small intestine from naïve or infected mice (D7), per crypt **(J-P)** Bulk RNA sequencing of 1 the first centimeter of duodenum in wildlings or SPF mice 7 days post-infection with 500 L3 of *N. brasiliensis* (n=4). **(J)** Volcano plot representing DEGs affected by wildlings as compared to SPF. **(K&L)** Over-representation analysis showing wildlings upregulated (K) or downregulated (L) DEGs, performed in Metascape (DEGs selected at p=0.05). **(M)** Heatmap of manually curated genes associated with tuft cell identity or activation/regulation, expressed as z-score. **(N)** Enrichment plots generated by GSEA analysis of tuft cells as defined in Habes et al **(O)** Enrichment plots generated by GSEA analysis of tuft cells subspopulation as defined in Buissant des Amories et al **(P)** SPF and wildlings small intestinal crypts were culture with or without Wnt3 supplement for 6 days. **(Q)** Enrichment plots generated by GSEA analysis of the Wnt signalling pathway.

Under succinate treatment, SPF mice displayed the expected infection-induced goblet cell hyperplasia, whereas wildlings again failed to mount a robust early response **(Fig. 4D)**. Goblet cell mucus content increased with infection in both groups; however, wildlings maintained significantly smaller goblet cells compared to SPF counterparts **(Fig. 4E)**. In line with the low tuft cells and goblet numbers in wildlings, ILC2s were also markedly constrained in wildlings **(Fig. 4F)**. Thus, augmenting luminal sensing was insufficient to restore epithelial effector function in microbiome-experienced mice.

The inability of succinate to rescue tuft expansion suggested a broad intrinsic hypo-responsiveness. We therefore asked whether epithelial responses to type 2 cytokines are intrinsically altered by microbial experience, independently of cytokine availability. To test this directly, naïve mice were supplemented with succinate or treated with IL-13/Ab(IL-13) complex prior to tissue collection **(Fig. 4G, Fig S4A)**. In SPF mice, both succinate and IL-13 induced robust tuft cell hyperplasia relative to untreated controls **(Fig. 4G, Fig S4A)**. In contrast, wildlings exhibited no significant increase in tuft cell numbers under either condition. Strikingly, goblet cells were increased by IL-13 treatment in SPF as well as in Wildling mice **(Fig 4H)**. In mice, tuft cell expansion in response to IL-13 stimulation is thought to arise from Lgr5+ stem cell commitment towards the secretory lineage^24^. Our results suggest that Lgr5+ stem cells are still responsive to IL-13 as goblet cells can expand in wildlings upon this stimuli, suggesting the differentiation block would be at the early progenitor stage of tuft cells (Tuft-P cells). Those Tuft-P cells have been reported to be present in crypts or in the crypt/villi junction^24^. Analysis of crypt-resident DCLK1 cells revealed that infection induced early tuft lineage commitment in both mice type. However, there was still significantly less cells in the wildlings after infection (**Fig 4I**).

To define the molecular basis of this hypo-responsiveness, we performed bulk RNA sequencing of small intestinal tissue at day 7 post-infection. Principal coordinate analysis revealed clear segregation between wildlings and SPF samples **(Fig. S4B)**. Differential expression analysis identified 329 upregulated and 446 downregulated genes in wildlings **(Fig. 4J)**. Upregulated pathways in wildlings were enriched for adaptive responses to commensals and pathogens, with strong Th1- and Th17-associated inflammatory signatures, as well as enhanced neutrophil “degranulation” and “chemotaxis” pathways **(Fig. 4K)**. We further confirmed enhanced level of IFNg but not IL-17 in intestinal tissue by MSD, IL-10 was also not found different **(Fig. 3N)**. In contrast, downregulated pathways were predominantly associated with epithelial and neuromuscular functions, consistent with a delay in the weep and sweep response **(Fig. 4L)**. In line with our histological observation, pathways linked to tuft cell activation, including “c-Kit receptor signaling” and “PLC-β–mediated events,” were reduced in wildlings. c-Kit expression was recently shown to be essential specifically for IL-13-induced tuft cell hyperplasia^25^, while remaining dispensable for baseline homeostatic maintenance. This mirrors the phenotype observed in wildlings and suggests a selective defect in the expansion of tuft cell progenitors.

To further define potential constraints on the development of intestinal epithelial cells after Nb infection in wildlings, we performed GSEA analysis with intestinal epithelial cells subsets as defined by Haber et al^26^. Enterocytes and enterocrine cells were found to be similar between wildlings and SPF mice **(Fig S4C).** Wildlings showed a trend toward increased goblet cell gene expression **(Fig S4C)**, despite lower number of those cells observed by histology. In line with the DEGs **(Fig 4M)** and our histological analysis, the tuft cell gene set was found enriched in SPF mice **(Fig 4N)**.

Recently, tuft cells have been shown to differentiate from Lgr5+ cells into tuft-p cells, then they further differentiate into tuft-1 and tuft-2 cells in a gradation along the crypt to villi axis^24^. To further characterize the wildlings tuft cells hypo-responsiveness, we asked if one of those subsets of tuft cells were particularly affected. We again performed GSEA analysis of our bulk RNA sequencing using gene sets for tuft-p, tuft-1 and tuft-2 defined from Buissant des Amorie et al^24^. Tuft-2 and Tuft-1 gene set were found significantly increased in SPF mice **(Fig 4O)**, in line with the low number of tuft cells found in the villi by DCLK-1 staining. Tuft-p analysis showed an interesting pattern, with some genes positively correlated with SPF mice and some negatively correlated with wildlings, suggesting a potential block in the differentiation of tuft-p cells in wildlings **(Fig 4O)**. Leading genes, strongly enriched in SPF mice and with low expression in wildlings, include Lpar3, Kcnmb1, Kcnab1, Aqp4 in line with the chemosensory role of tuft cells, Ugt2a3, Ugt2b36, Ces1d, Akr1c14, Hsd3b3 linked to metabolic and detoxification activity, as well as 3 transcription factos namely Foxa2 (Forkhead 2, a pioneer transcription factor known to be involved in epithelial differentiation and secretory lineage competence), Hes2 (a member of the HES family of Notch repressors), Nr2e3 (an orphan nuclear receptor), as well as two regulatory genes Zfp667 and Samd5 **(Fig S4D)**.

To determine whether the hypo-responsiveness of Wildling tuft cells to IL-13 is intrinsic or extrinsic, we generated small intestinal organoids. Strikingly, Wildling-derived organoids exhibited slower growth than those from SPF mice, with fewer buds observed six days after the start of culture. In human intestinal organoids, Wnt supplementation is required to maintain stem cell proliferation^27^. Accordingly, the addition of Wnt to Wildling organoid cultures supported growth and budding, restoring a pattern similar to that observed in SPF mice **(Fig. 4P).** To confirm wildlings have a “Wnt-low” signature, we performed GSEA on Wnt signaling pathway(mmu04310). We confirmed Wnt signaling is enriched in SPF mice relative to Wildlings **(Fig. 4Q)**. Leading-edge analysis of the suppressed Wnt signature in Wildlings revealed a multi-level collapse of the pathway. This included a significant reduction in essential crypt-base ligands (Wnt3, Wnt2b) and their associated receptors (Lrp5/6, Fzd family), suggesting a diminished signalling capacity of the Wildling intestinal niche. Crucially, we observed the concurrent downregulation of a suite of transcriptional co-factors and chromatin remodelers, including Chd8, Tbl1x/xr1, Ep300 and Crebbp, the last two being required for the histone acetylation (H3K27ac) that licenses Kit expression.

Collectively, these data suggest that a microbiome dependent Wnt-tune down in the Wildling crypt creates a chromatin-level bottleneck at the Kit locus, selectively blunting Tuft cell progenitor expansion while sparing Notch-dependent lineages like goblet cells.

### Subhead 5: Fecal microbiota transfer selectively recapitulates epithelial tuft hypo-responsiveness but not systemic type 2 suppression

To determine whether the delayed expulsion and suppressed type 2 responses observed in wildlings reflect early-life immune conditioning or ongoing microbial exposure, we performed fecal microbiota transfer (FMT) into adult SPF mice. Eight-week-old SPF mice were colonized with cecal contents from wildlings (W→SPF) or SPF controls (S→SPF), while a third group remained ungavaged. Following three transfers administered every other day and a 20-day colonization period, mice were infected with *N. brasiliensis* **(Fig. 5A)**.

**Fig. 5.**
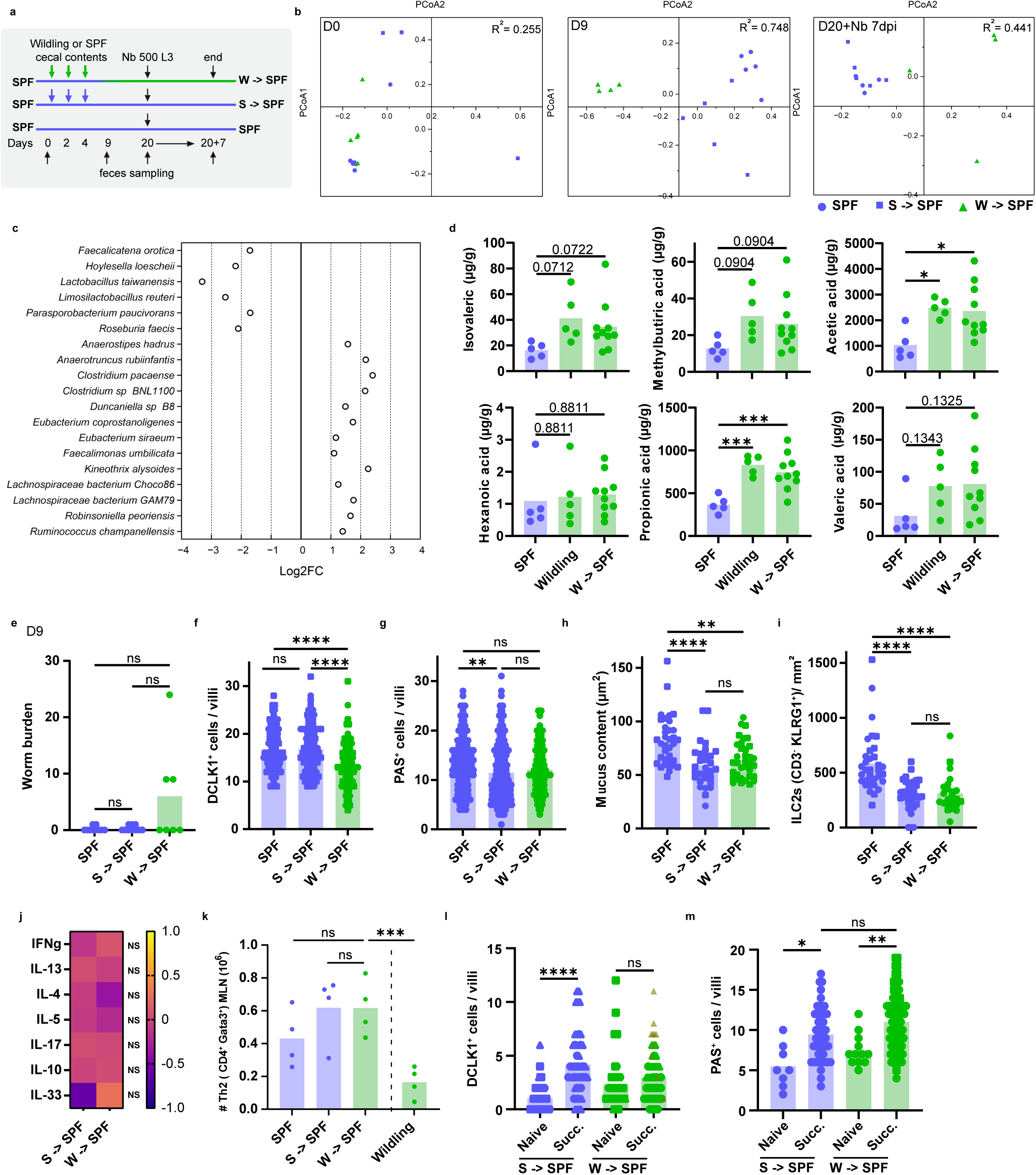
Natural microbiome from wildlings transferred at adult age recapitulate the dampened of the “weep and sweep” but not of “Th2 response”. **(A-M)** Caecal material from wildlings or SPF was transplanted by oral gavage into SPF mice. One SPF group was left untreated. After 20 days post colonization, all mice were infected with 500 L3 of *N. brasiliensis* for 7-9 days. Fecal pellets from all mice were collected over 27-29 days and long read16S rRNA gene profiling by nanopore was performed. **(B)** PCA analysis of microbial 16S sequencing in mice described in A (n=4-5 mice) at 0, 9 and 20 days post colonization. **(C)** Differential abundance of bacterial taxa at day 20 post-colonization, shown as log2 fold change (Log2FC); negative values indicate enrichment in W→SPF mice, positive values enrichment in SPF controls. **(D)** Fecal metabolites measured by non-targeted metabolomics**. (E)** Worm burden was measured by baermann in the intestine. **(F)** Tuft cell quantification by immunofluorescence analysis of DCLK1 in the small intestine from naïve or infected mice, per villi. **(G&H)** PAS staining of Swiss-roll of 3 anterior cm of small intestine. Quantification of the number of Goblet cells per villi (G) or of the area of Goblet cells (H). **(I)** Intestinal ILC2 number quantified by immunohistology (CD3- KLRG1+) **(J)** Intestinal cytokines measured by MSD in infected mice in SPF or wildlings for 7 days. Pool of 2 experiments (n=3-6 mice). **(K)** Number of Th2 cells (I, CD45^+^ CD4^+^ Gata-3^+^) in the MLN of colonized mice. Representative of 2 independent experiments, (n=4-6) **(L-M)** FMT mice as described in A were stimulated for 7 days with succinate in the drinking water **(L)** Tuft cell quantification by immunofluorescence analysis of DCLK1 in the small intestine from naïve or infected mice, per villi. **(M)** PAS staining of Swiss-roll of 3 anterior cm of small intestine. Quantification of the number of Goblet cells per villi. When possible, different symbols are used between biological repeats.

To determine whether fecal transfer established a microbial ecosystem associated with tuft cell hypo-responsiveness, we performed longitudinal 16S rRNA Nanopore sequencing on fecal samples collected before, during, and after colonization **(Fig. 5A)**. Prior to gavage, all groups displayed comparable microbial composition **(Fig. 5B)**. Principal coordinate analysis revealed clear segregation of W→SPF mice at both day 9 and day 20, confirming stable establishment of a distinct microbial community **(Fig. 5B, FigS5a)**. By day 9 following the first transfer, W→SPF mice exhibited a marked compositional shift characterized by enrichment of Firmicutes, including Lactobacillaceae, and introduction of Bacteroidetes families such as Prevotellaceae and Bacteroidaceae. By day 20 post-colonization, this shift consolidated with expansion of Clostridiales, Selenomonadales, and Bacteroidales **(FigS5a)**. These compositional changes were accompanied by a significant increase in alpha diversity in W→SPF mice compared to controls at day 20 **(Fig. S5C)**. Of note, the engraftment was not successful for one mouse that segregated with SPF mice and showed low egg burden **(FigS5D),** indicating high variability with the FMT approach. Consistent with this restructuring, differential abundance analysis highlighted a reciprocal enrichment pattern, with several Lactobacillaceae and other taxa (e.g., *Lactobacillus taiwanensis*, *Limosilactobacillus reuteri*) preferentially associated with Wildling-derived microbiota, while multiple Clostridiales and Lachnospiraceae members were enriched in SPF-recipient communities; functionally, this compositional shift correlated with a remodeling of microbial metabolic output, including reduced predicted butyrate/isobutyrate production capacity in W→SPF mice compared to SPF controls **(Fig. 5C).**

In line with this altered fermentative landscape and with previous analyses in naïve wildlings^19^, targeted metabolomic analysis revealed that W→SPF mice recapitulated the wildling-associated increase in acetate and propionate levels, confirming the establishment of a distinct, wildling-like metabolic niche **(Fig. 5D).**

Despite this change in microbiome, colonization with wildling microbiota was not sufficient to reproduce the delayed expulsion phenotype. Although a subset of W→SPF mice retained worms at day 7 post-infection, overall parasite clearance did not significantly differ from SPF or S→SPF controls **(Fig. 5E)**. Thus, adult microbiota transfer alone does not recapitulate the impaired expulsion observed in wildlings.

Despite the absence of significant delayed clearance, W→SPF mice exhibited significantly reduced tuft cell numbers following infection compared to both SPF and S→SPF controls **(Fig. 5F)**, indicating that tuft hypo-responsiveness is at least partially transferable in adulthood. Both goblet cell numbers and their mucus content were not different in W→SPF as compared to S→SPF controls **(Fig. 5G&H),** suggesting the tuft cells activation level might still be enough to activate the tuft/ILC2 circuit in W→SPF mice. Similarly, ILC2 density was decreased as compared to SPF mice but not different between W→SPF and S→SPF mice, suggesting that this effect reflects a general consequence of cecal gavage rather than a specific feature of the wildling microbiota **(Fig. 5I)**.

We previously observed elevated intestinal IFN-γ and reduced IL-4 and IL-5 levels in wildlings during infection, consistent with a type 1-skewed environment. These cytokine alterations were not reproduced in W→SPF mice, as compared to S→SPF mice **(Fig. 5J)**. IFN-γ levels were comparable across groups, and reductions in IL-4 and IL-5 were observed in both W→SPF and S→SPF mice, in line with the generalized decrease in ILC2s. These data indicate that the inflammatory skewing characteristic of wildlings is not transferred by adult microbiota colonization.

Consistent with this distinction, mesenteric lymph node Th2 responses were preserved in W→SPF mice. Th2 cell numbers did not differ significantly between W→SPF, S→SPF, and SPF controls **(Fig. 5K)**, and remained substantially higher than those observed in wildlings. Thus, systemic suppression of type 2 immunity is not recapitulated by fecal transfer.

To directly test whether tuft cell hypo-responsiveness was selectively transferred during FMT, we assessed epithelial responsiveness to succinate supplementation. As observed in SPF mice, succinate induced tuft cell expansion in S→SPF controls **(Fig. 5I)**. In contrast, W→SPF mice failed to mount a tuft cell expansion in response to succinate, confirming that impaired metabolite-driven activation of tuft cells is transferable with the wildling microbiota. Like for the Nb infection, W→SPF mice mounted a strong goblet cells response to succinate treatment similar to S→SPF controls, confirming the constraint is specifically on tuft cells, downstream of the common secretory lineage progenitor commitment.

## DISCUSSION

Soil-transmitted helminths establish chronic intestinal infections in humans within ecologically complex environments shaped by lifelong microbial and pathogen exposure. In contrast, most mechanistic insights into anti-helminth immunity derive from specific pathogen-free (SPF) mouse models that develop in the absence of such ecological conditioning. Using a comparative approach between SPF mice and wildlings harboring a vertically transmitted, complex microbiota^1,20^, we find that parasite establishment proceeds similarly across housing conditions, but the magnitude and organization of type 2 immunity differ markedly. Wildlings mount a controlled and tissue-protective response characterized by limited pulmonary damage, restrained ILC2 expansion, and macrophage polarization, whereas SPF mice exhibit amplified type 2 cytokine production and more pronounced tissue pathology. In the intestine, wildlings display a calibrated epithelial program with reduced tuft cell amplification and delayed worm expulsion, while SPF mice generate the robust “weep and sweep” response classically described in this model. Importantly, epithelial calibration could be partially transferred to adult SPF recipients by fecal microbiota transfer, whereas systemic type 2 amplification remained a feature of SPF developmental history, indicating early-life imprinting of lymphoid compartments. Finally, epithelial changes in wildlings associated with enrichment of fermentative bacterial taxa and elevated acetate and propionate levels, linking ecological microbial complexity to barrier immune setpoints. Together these findings support a model in which early-life microbial experience establishes a physiologic immune equilibrium that constrains excessive type 2 amplification while preserving protective capacity.

Although our study focuses on the experimental STH model *Nippostrongylus brasiliensis*, which induces an acute and self-limiting infection in laboratory mice, human soil-transmitted helminths typically establish chronic infections within ecologically complex hosts^28,29^. The robust “weep and sweep” response described in SPF systems may therefore represent one extreme of a broader physiological spectrum. The phenotype observed in wildlings aligns with emerging data from ecologically exposed systems. Rewilding studies in laboratory mice have demonstrated altered expulsion kinetics and decrease type-2 response following *Trichuris muris* infection^18,30^, while field studies in wild wood mice indicate that chronic *Heligmosomoides polygyrus* infection persists despite measurable immune activation, including type 2-associated responses, and is characterized by marked immune heterogeneity and limited correlation between canonical Th2 markers and parasite clearance^31,32^. These models collectively indicate that helminth immunity in natural settings operates under constraints distinct from those observed in SPF conditions. Wildlings provide a complementary platform that bridges ecological realism and experimental control: unlike rewilded mice, microbial and pathogen exposure is vertically transmitted from birth, and unlike wild-caught rodents, wildlings retain the genetic background and tractability of laboratory strains. This combination permits dissection of how lifelong ecological conditioning shapes epithelial-immune circuits while preserving the capacity for controlled infection, transfer, and mechanistic intervention.

A central paradigm in anti-STH immunity is the intestinal tuft cell-ILC2 circuit^22^, in which parasite metabolites trigger tuft cell release of IL-25, activating ILC2s to produce IL-13 and drive expansion of both epithelial tuft and goblet cells. This “weep and sweep” mechanism has been well characterized in SPF models and is classically thought to be required for rapid parasite expulsion. In wildlings, restraint occurred at multiple levels of this circuit, including limited tuft cell expansion and altered transcriptional programs in intestinal tissue, decreased ILC2 and goblet cell expansion. The incomplete and variable transfer of the tuft phenotype by FMT suggests that multiple ecological layers regulate tuft lineage amplification, potentially including non-bacterial microbial components and developmental imprinting.

Microbiota-derived metabolites have recently emerged as regulators of epithelial differentiation, with butyrate shown to restrict tuft cell hyperplasia via HDAC3-dependent mechanisms^33,34^. Although butyrate was not significantly elevated in our system, acetate and propionate were enriched in wildlings and transferable to SPF recipients of wildling caecal material. These metabolites are known modulators of epithelial and immune cell function, raising the possibility that fermentative microbial communities, that correlated with low tuft cell response, tune epithelial activation thresholds. However, direct causality between specific metabolites and tuft cell restraint remains to be established. Of note, butyrate treatment and HDAC dependent mechanism affect tuft cells differentiation under baseline conditions, suggesting a general alteration of tuft cell lineage commitment. In wildlings, tuft cell numbers are reduced only in the context of type 2 response and not at baseline, highlighting a marked difference between those systems. To our knowledge the only reported defect affecting similarly IL-13 induced tuft cells rather than baseline tuft cell number is a deficiency in KIT, that affects early commitment of tuft cells under type 2 conditions, downstream of Pou2f3 expression^25^. Interestingly, KIT was notably reduced in our bulk RNA sequencing and the early commitment block is in line with our observed altered transcriptional profile of tuft-p cells in wildlings. Of note, KIT deficiency or treatment with Imatinib reduced tuft cells expansion after *N. brasiliensis* infection^25^ but not to the extent observed in wildlings, suggesting other mechanisms are likely at play under natural microbial conditions.

ILC2 expansion in wildlings caecal transfer recipients occurred despite limited tuft cell amplification, suggesting that alternative activation pathways may sustain type 2 cytokine production under ecologically conditioned settings or that a certain threshold of tuft cells is sufficient to ensure efficient downstream effector mechanisms. ILC2s integrate neuronal, stromal, and lipid-derived signals *in vivo*, many of which are known to be influenced by microbial composition. Moreover, we observed preservation of IL-13 levels in the intestine despite reduced ILC2 and Th2 cell frequencies and decreased IL-5 and IL-4 levels in wildlings, indicating that epithelial restraint does not necessarily equate to global cytokine deficiency under natural conditions. This dissociation suggests altered cellular contributions or spatial organization of IL-13 production that warrants further investigation.

Overall, our data delineates three layers of immune calibration imposed by microbial experience: a transferable epithelial restraint associated with fermentative microbial ecosystems, and two developmentally imprinted systemic tone that constrain tuft cell lineage commitment under IL-13 stimulation on one hand and limits excessive type-2 amplification on the other hand. Rather than representing immune deficiency, this state reflects a physiological equilibrium shaped by ecological exposure which mitigates tissue damage while maintaining the capacity for parasite clearance. These findings underscore the importance of considering microbial and developmental context when defining canonical immune mechanisms and suggest that the intensity of type-2 amplification observed in SPF systems may represent only one end of the adaptive landscape of helminth immunity.

## MATERIALS AND METHODS

### Study design

The goal of this study was to define the impact of ecological realism on the development of protective type-2 responses against helminths. To address this, we used a comparative approach between clean SPF C57BL/6TacN mice and wildlings and characterize the lung and intestinal immune response after Nb infection using histology, flow cytometry, and scRNA-seq, together with microbiome sequencing and metabolomics. Sample sizes were based on prior experience for SPF mice and had to be re-assessed for wildlings with every new data generation due to higher variability between those mice. All key findings were confirmed in two to three independent experiments using multiple biological replicates (3-8 per experiment). Animals were randomly assigned to experimental groups based on genotype, sex, age and availability, and data were analyzed using predefined gating strategies. All samples meeting predefined quality criteria were included, and data were excluded only in cases of clear technical failure. The number of mice per group, the number of experimental replicates, and the statistical tests used are reported in the figure legends. All data points are biological replicates or in the case of intestinal histological analysis are a given villi/crypts to fully capture the diversity of response in the tissue.

### Mice and hygiene standards

SPF control mice were C57BL/6NTac mice purchased from Taconic Biosciences, Denmark. wildling mice were bred on a C57BL/6NTac background and were provided by S. Rosshart from two separate facilities in Uniklinik Erlangen or Uni Freiburg (Germany); key results were stable and repeatable in animals from both facilities. As previously reported, wildling colonies were generated by embryo transplant into pseudo-pregnant wild mice captured on farms, and the subsequent generations of wildlings were supplemented with soil and grass. Age- and sex-matched mice (8 to 16 weeks old) were used. Animals were housed in a conventional facility but in individually ventilated cages, facilitating the maintenance of SPF status. A 12/12 hour light-dark cycle, at 20-24°C, and 45-65% humidity is maintained in the facility. Water and standard diet were provided non autoclaved *ad libitum*. Environmental enrichment (*e.g.* nesting material, tunnels) was provided in all cages and all procedures complied with institutional and cantonal welfare regulations (BL543 and BL560). Euthanasia was performed by ketamine/xylazine overdose. wildling and SPF mice were housed in the same room of the animal facility. Husbandry followed strict quarantine procedures, inside a biosafety cabinet and using Bacillol AF for disinfection. FELASA testing was performed according to IDEXX BioAnalytics guidelines (blood on Opti-Spot Cards, fur swabs, fecal pellets). Mouse 3R FELASA Annual (Serology) and Mouse 3R FELASA Annual (PCR) panels were conducted to ensure the SPF status of mice in comparison to wildlings (in 6 independent cohorts of mice, across 4 years of experiments).

Breeding pairs used to generate wildlings for our experiments were treated with fenbendazole-supplemented chow to remove pinworms. In compliance with facility quarantine guidelines, wildlings were given fenbendazole for one week before entering the Swiss TPH animal facility. During a five-day acclimatizing period, wildling and SPF control mice were given fenbendazole-supplemented chow to control for any effect of treatment. A minimum of 3 cage changes and one week was allowed to pass after fenbendazole treatment before infection with *N. brasiliensis* L3.

### Parasite lifecycle and infections

*N. brasiliensis* was maintained as previously reported^35^ In brief, Lewis rats were infected with 3000-5000 third-stage larvae (L3) (the Prof. Lindsay Dent strain kindly provided by Prof. Graham LeGros, MIMR, New Zealand at installation at the Swiss TPH). In the evening of day 5 to day 8 post-infection, rats were gridded overnight, and feces used for coproculture. iL3 were collected from the edge of coproculture plates and washed extensively in PBS before subcutaneous infection with 500 L3 in 100uL in the neck using a 25G needle and 0.5mL insulin syringe.

### *In vivo* treatments

IL-13-anti-IL-13 complexes were prepared by mixing 5 µg of recombinant mouse IL-13 (Peprotech, Gibco, Cat. # 210-13-10UG) with 25 µg of IL-13 monoclonal antibody (eBio1316H, Cat. # 16-7135-85, Invitrogen) in PBS. IL-13-anti-IL-13 complexes or vehicle (PBS) was injected intraperitoneally every 2 days and culled 4 days after the start of treatment.

Mice were given autoclaved drinking water supplemented with 150 mM sodium succinate hexahydrate ad libitum for 7 days in black bottles.

The ileocecal microbial communities from terminal ileum and cecum of wildlings and SPF mice were transferred by a three oral gavages every other days before at least 20 days of recolonization. Recipient mice were C57BL6Ntac 8 weeks old male and did not receive any treatment prior microbial engraftment. The caecal extract was collected as described^19^. Briefly, the caecum were collected in a biosafety cabinet class 2 with autoclaved tools and placed in a sterile bacteriological petri dish without opening. The caecal content was extruded by squizzing with tweezers into a sterile 100 µm cell strainer (Corning Falcon cat. #352360) placed inside of a sterile bacteriological petri dish. The collected fecal material was weighed and then diluted at a 1:2 ratio (g collected material: ml fluid) in sterile filtered cryoprotectant (PBS (Panbiotech, P04-361000) containing 0.1% L-cysteine (Sigma-Aldrich, C7352) and 20% glycerol (Sigma-Aldrich, G2025)). The resulting suspension was subsequently mashed through the 100 µm cell strainer to remove all insoluble particles, 800 ul aliquots were prepared into 2 mL externally threaded plastic screw-capped cryo-vials (Azenta Life Sciences, BCS-2502), subjected to controlled freezing with 1 °C per minute (Azenta Life Sciences CoolCell LX Cell, BCS-405) and stored in a −80 °C mechanical freezer. 100 ul of caecal material was inoculated to each mouse per gavage.

### Enumeration of worm burden

Worm burdens and fecal egg count were determined as previously described^35^. If required, broncho-alveolar lavage (BAL) was performed with 3 times 1 mL cold PBS. Lungs were removed from infected mice and minced and placed in a baerman apparatus. Larvae isolated from Baerman or filtered from BALF were counted using a dissection microscope.

For intestinal worm burden, the number of worms on tissues that required specific processing (histology or RNA) were directly counted after opening the intestinal segment on a stereomicroscope. The rest of the tissue was processed by baerman as for the lungs.

For caecal egg count, caecal content was squeezed out of the caecum, weighed, and dissolved in aqueous saturated NaCl. Buoyant eggs were counted using a McMaster counting chamber and normalized to fecal weight.

### Preparation of single-cell suspensions from lungs

Bronchoalveolar lavage was performed by three consecutive washes of 1 mL of cold Dulbecco’s phosphate buffered saline (PBS) (Bioconcept, 3-05F00-I) using the catheter of an 18-G Vialon needle (BD Insyte Catheter, 381444). The first 1mL of BAL was centrifuged and the supernatant taken for cytokine analysis. The cell pellet was pooled with the remaining BAL, centrifuged, and the pellet counted for red blood cell counts to measure hemorrhage. The remainder of the cell pellet was processed by incubating with 3mL of red blood lysis buffer (Roche, 11814389001) at room temperature or 37°C for 5 min depending on the abundance of hemorrhaging.

Lungs were resected and immediately placed in DMEM (Sigma-Aldrich) supplemented with 10% fetal calf serum (Bioconcept, 2-01F00-I) on ice. In a sterile biosafety cabinet, lungs were minced in dry 6-well plates then 3mL of freshly prepared digestion buffer was added, 0.48mg/mL collagenase type 1 (Gibco, 7100017) and 500U/mL DNase 1 recombinant (Roche, 04536282001) in RPMI (Bioconcept, 1-41F51-I). Plates were placed in a pre-warmed orbital shaker at 170rpm at 37°C for 45 minutes. The reaction was stopped on ice, aspirated (approx. 10 times) with an 18-G needle and 5mL syringe and passed through a 70um cell strainer (Corning, 431751) on a 50mL falcon tube (Corning). Lung tissue was crushed with the end of a 1mL syringe plunger with an additional 10mL of DMEM, 10% FCS. Cells were centrifuged at 400 x *g*, 5 min 4°C, and red blood cell lysis (Roche, 11814389001) performed at room temperature. For tissues harvested at the peak of hemorrhaging (day 2 post-infection) red blood cell lysis was performed at 37°C, 5% CO2. Cells were counted using Trypan blue.

### Preparation of single-cell suspensions from mLN

Mesenteric lymph nodes were dissected, taking care to remove the entire chain including the duodenum draining node. The chain was transferred to PBS, 10% FCS on ice. In a sterile biosafety cabinet, fat tissue was removed and placed on sterile glass slides with graphing paper underneath and photographs taken for area measurements in FIJI ImageJ. Single-cell suspensions were prepared from MLNs by placing on a moistened 70uM cell strainer inside a 5mL petri-dish and crushing the node with cold PBS, 2%-10% FCS, using the handle of a 1mL syringe. Cells were centrifuged at 400 x *g* for 5 min, 4°C, and the cell concentration was counted by trypan blue.

### Intestinal histology

The small intestinal length was measured using a ruler. 2-3 cm of small intestine were collected for swiss roll preparation. In general, the first 3 cm were used for paraffin sectioning and the 3 next ones for OCT sectioning.

Small intestinal tissue morphology was studied using the ‘Swiss rolling’ technique. For paraffin embedded Swiss rolls, 3cm of the duodenum was removed (beginning from the second cm of the intestine) and gently flushed with 2mL of cold PBS using a 18-G Vialon catheter (BD Insyte Catheter, 381444). The intestine was then inverted on a toothpick pre-soaked in PBS and immediately placed in ice cold methanol-free 4% paraformaldehyde (PFA) with 5% glacial acetic acid in PBS in a 15mL falcon tube. Inverted intestines were fixed initially at 4°C for 4 hours, then cut from the skewer, rolled and fixed again in the same tube overnight at 4°C before washing into 70% histology-grade ethanol and paraffin embedding. For OCT-embedded Swiss rolls, 3cm of intestine (beginning from the 5^th^ cm of the duodenum) was fixed in a similar manner in 4% PFA (no acetic acid), for a maximum of 4 hours at 4°C. After rolling, Swiss rolls were transferred to 15% sucrose in PBS for 8-24 hours, then 30% sucrose for 12 hours, and a 30% sucrose, 50% optimal cutting temperature medium (OCT) solution overnight at 4°C. Swiss rolls were cryoembedded on a dry ice-isopropanol bath.

Swiss rolls were sectioned at 6um. Antigen retrieval, after paraffin dewaxing in Histoclear and rehydration, was performed in pH9.0 Tris EDTA buffer, and sections were blocked and stained with rabbit polyclonal DCLK1 (Abcam, anti-DCAMKL1 ab31704, 1:200) and donkey anti-rabbit highly cross adsorbed AlexaFluor 568 secondary antibody (Invitrogen, A10042, 1:500). For ILC2 staining, OCT sections were used and stained after permeabilization in PBS, 1%(v/v) donkey serum, 0.1 % triton-X for 10 min, with anti-CD3 (clone 17A2, biolegend, RRID AB_312658, 1:50) for 1 hour and anti-KLRG-1 (clone 2F1, eBioscience, RRID AB_469131, 1:50) overnight. Goblet cell (PAS) staining was performed after dewaxing using 1% periodic acid solution (10 mins), Schiff reagent (15 mins) followed by Mayer’s hematoxylin and Scott’s blueing reagent counterstaining. Slides were mounted in Eukitt mounting media (Sigma-Aldrich, 03989). PAS and immunofluorescence slides were imaged on a Zeiss Axio Scan.Z1 slide scanner at 20 (tuft cells and PAS staining) or 40X (ILC2s).

Cells were counted manually after visualization in FIJI Image J (v 1.54f). Results are plotted per villi or crypt or per analysed villi surface for ILC2s. For goblet cell area measurements, cells were measured by manual thresholding and delimiting the cell limit. Investigators were blinded during histological quantification.

### Flow cytometry

Isolated lung cells were plated into round-bottom plates (Thermo Scientific, 10344311) and 1×10^6 cells (or 2×10^6 for ILC staining) were washed into PBS. LIVE/DEAD Fixable Violet Dead Cell (Invitrogen, L34955) staining was performed (1:250) on ice, 20min, then cells were incubated with Fc block (CD16/CD32, BioXCell clone 2.4G2, BE0307) followed by a relevant staining mix. BAL fluid was stained with CD45-BV510 (clone 30F11, Biolegend, 1:100), Ly6G-AF647 (clone 1A8, Biolegend, 1:200), Gr1-FITC (clone RB6-8C5, Biolegend, 1:100), CD11b-PerCP (clone M1/70, Biolegend, 1:100), SiglecF-PE-Dazzle CF594 (clone S17007L, Biolegend, 1:200), I-A/I-E (MHC-II)-APC/Fire750 (clone M5/114.15.2, Biolegend, 1:100). Neutrophils were identified as CD4^5+^ Ly6G^+^ Gr1^+^ cells. Alveolar macrophages were identified as CD45^+^ LyG-SiglecF^+^ I-A/I-E^+^ cells and Eosinophils as CD45^+^ LyG-SiglecF^+^ I-A/I-E- SCC^high^ cells.

For ILC2 and Th2 cell staining, cells were stained with CD45-BV510 (clone 30F11, Biolegend, 1:100), CD25-BV785 (clone PC61, Biolegend, 1:400), CD4-APC/Fire750 (clone GK1.5, Biolegend, 1:400), KLRG1-BV605 (clone 2F1, Biolegend, 1:200), CD127-AF700 (clone A7R34, Biotechne/R&D, 1:100), FceRIa-PE-Cy7 (clone MAR-1, Biolegend, 1:400), and c-kit-PerCP (clone 2B8, Biolegend, 1:100). A lineage dump channel was used for ILC identification: SiglecF-PE (clone S17007L, Biolegend, 1:200), Gr-1-PE (clone RB6-8C5, Biolegend, 1:200), B220-PE (clone RA3-6B2, Biolegend, 1:200), CD11b-PE (clone M1/70, Biolegend, 1:200), CD11c-PE (clone N418, Biolegend, 1:150), NKp46-PE (clone 29A1.4, Biolegend, 1:100), CD8-PE (clone 53-6.7, Biolegend, 1:400), and F4/80-PE (clone BM8, Biolegend, 1:400). The eBioscience Foxp3/Transcription Factor Staining buffer set (Invitrogen, 00-5523-00) was performed before overnight intracellular staining with GATA3-AF488 (clone TWAJ, Invitrogen, 1:100), and FoxP3-APC (clone FJK-16s, eBioscience, 1:100). ILC2s were identified as Lin^−^KLRG1^+^ Gata-3^+^ cells. Th2 cells were identified as CD4^+^ Gata-3^+^ cells.

For identifying macrophage populations in the lung, we stained with CD45-BV510 (clone 30F11, Biolegend, 1:400), F4/80-APCeF780 (clone BM8, eBioscience, 1:200), CD11c-BV785 (clone N418, Biolegend, 1:200), SiglecF-PE/Dazzle594 (clone S17007L, Biolegend, 1:200). Cells were permeabilized with ice-cold methanol, rehydrated and stained overnight with RELMa-AF700 (clone DS8RELM, eBioscience, 1:100), Phospho-STAT6-PercP eFluor710 (clone CHI2S4N, eBioscience, 1:100), and Arg1-APC (Polyclonal #P05089, R&D, 1:100). Alveolar macrophages were identified as CD45^+^ F4/80^+^ Cd11c^+^ cells. Interstitial macrophages were identified as CD45^+^ F4/80^+^ AM^−^ SiglecF^−^ cells. Eosinophils were identified as CD45^+^ F4/80^+^ AM^−^ SiglecF^+^ cells.

Cells were re-fixed with 2% PFA or eBioscience Foxp3 buffer and acquired on a Beckman Coulter CytoFlex S. Data was analysed in FlowJo v.10.6.1 and bar plots made in GraphPad Prism 10.

### ELISA and MSD

1 cm intestinal tissue was snap-frozen at collection and lysed in 200 ul of Tris-lysis buffer containing protease inhibitors with 3 2.3 mm diameter Zirconia/Silica beads (Bio spec products, 11079125Z) on the “mouse intestine” program on a FastPrep®-24 5G bead beating grinder and lysis system (116005500). Lung tissue was homogenized in 100ul PBS containing protease inhibitors as described for the intestine. Both tissues were centrifuged after homogeization at 11000 x g, 15 min, 4C and supernatant was frozen until use for ELISA or MSD detection.

Cytokines were assessed in lung and intestine homogenate using a MSD U-plex Group1 kit according to the manufacturers’ instructions and measured using a MESO QuickPlex SQ120. Lung cytokine levels were standardized to tissue weight.

### RNA extraction and bulk sequencing

For the intestine, the inflamed 1^st^ cm of the duodenum was carefully processed to preserve RNA integrity. Without exposing the lumen, immediately upon confirming euthanasia, the intestine was exposed, and 1 cm of intestine was removed and submerged in cold RNALater mixed by inversion and placed on ice. Samples were stored overnight at 4°C and processed the next day two-at-a-time to minimize RNA degradation. Excess RNALater was removed and the tissue was placed in TRIzol (Invitrogen) and homogenized using three 2.0mm zirconia beads (Biospec, 11079124ZX) using the ‘mouse intestine’ program of a FastPrep-24™ 5G beater (MP biomedicals). Total RNA was extracted from tissue using the Qiagen AllPrep DNA/RNA Mini Kit (Cat. no. 80204) according to the manufacturer’s instructions. Integrity was assessed using the Agilent TapeStation RNA ScreenTape assay. Bulk RNA sequencing of small intestinal tissue was performed by Novogene using the Illumina NovaSeq X Plus platform (paired end 150 bp reads) following RNA sample QC, mRNA library preparation with poly(A) enrichment and standard data analysis workflows.

### Single-cell RNA Sequencing

For single-cell sequencing, BAL was not performed. Lungs were digested as for flow, and 10×10^6 cells per mouse were stained for the MACS Dead cell removal kit according to manufacturer’s instructions on LS columns. 0.5×10^6^ cells per mouse were then labelled for multiplexing with 10uL of sample tags from the BD® Mouse Immune Single-Cell Multiplexing Kit (633793) and 90uL of buffer according to manufacturer’s instructions (23-21340(02), 2022-11) in 1.5mL Eppendorf tubes pre-coated with BD® Stain Buffer (FBS) (554656). Samples were washed 3 times, resuspended in 300uL and counted in duplicate using Trypan blue and a hemocytometer. 100,000 cells per mouse from 5 SPF and 5 wildling mice were pooled into two respective groups, and 0.4×10^5 cells were used to over-load two BD Rhapsody cartridges according to the manufacturer’s protocol (23-21332(03), 2022-11) with BD Rhapsody™ Enhanced Cell Capture Beads v2.0. One cartridge was used for wildlings, and one for SPF mice. An Eppendorf ThermoMixer R5355. Cells were captured on the BD Rhapsody Express without Scanner. cDNA libraries were prepared using the BD Rhapsody Whole Transcriptome Analysis Amplification Kit (633801 BD Biosciences) following the BD Rhapsody System mRNA Whole Transcriptome Analysis (WTA) and Sample Tag Library Preparation Protocol (BD Biosciences). The final libraries were quantified using a Qubit Fluorometer with the Qubit dsDNA HS Kit (Q32851 Thermo Fisher Scientific). Library size distribution was measured with the Agilent high-sensitivity D5000 assay on a TapeStation 4200 system (5067-5592 Agilent Technologies). Sequencing was performed in paired-end mode (2 × 75 cycles) on a NovaSeq 6000 with NovaSeq 6000 SP Reagent Kit chemistry at ETH.

### Single cell sequencing analysis

Raw reads were processed with the BD Rhapsody Sequence Analysis Pipeline (version 2.1) using the Mus musculus reference genome (build GRCm39, gene annotation from GENCODE release M26). Read filtering, cell barcode / UMI extraction, alignment, distribution-based error correction, molecular identifier deduplication, cell calling (mRNA-based putative cell calling) and sample-tag based demultiplexing were performed with default pipeline parameters, retaining protein-coding, rRNA, tRNA, Mt rRNA and Mt tRNA transcript types.

Downstream analysis was performed on the resulting feature-barcode count matrices using the R package Seurat v5^36^. Per-sample quality control, filtering, normalization and clustering were performed using FGCZ’s standard ScSeurat workflow implemented in ezRun^37^. Cells were filtered according to UMI count, feature count, and mitochondrial percentage using a median-absolute-deviation (MAD) approach implemented in the package scater^38^. Cells with flagged as doublets by scDblFinder^39^ were removed. Filtered data were log-normalized and scaled using SCTransform implemented in Seurat^40^, using the top 3’000 highly variable genes. Dimensional reduction was performed using PCA. The Louvain algorithm was applied with a resolution of 0.6 to cluster the cells using the first 20 PCs.

Integration across all 10 samples was performed using the Harmony algorithm^41^ on SCTransform-normalized data, with batch correction based on biological Condition (Taconics vs. Wildlings). Clustering of the integrated dataset was performed with the Louvain algorithm (resolution 0.6) on the first 15 Harmony components. For each cluster, marker genes were identified by differential gene expression analysis (Wilcoxon rank-sum test with log2 fold-change > 0.25, adjusted p-value < 0.01, and a minimum fraction of cells expressing the marker > 0.1). Marker genes, combined with automated annotation against the ImmGen reference using SingleR^42^, reference databases provided through Enrichr^43–45^ and published literature were used for manual cell type annotation.

To resolve T-cell and ILC substructure, two lymphocyte subsets (Th2 / ILC2 / Treg compartment and a broader CD4/CD8 T-cell / ILC3 compartment) were extracted from the integrated object, re-normalized with SCTransform per sample, re-integrated with Harmony (correcting for Condition), and re-clustered on the first 10 Harmony components at a resolution of 1.2. Cluster annotation was refined using the same marker-based and reference-database approach as above.

Differential abundance and differential expression between Taconics and Wildlings were assessed per annotated cell type, using a pseudo-bulk approach (aggregated counts per sample and cell type) followed by Wilcoxon rank-sum testing. Sample-level similarity was visualized using a bulk-like PCA, in which expression was aggregated per sample with Seurat’s AggregateExpression, variable features were re-selected, data scaled, and the first two principal components were computed.

### Microbiome (16S) Sequencing

DNA was extracted from fecal (50mg) and mucus (1 cm of gut scraping, 5 to 70mg) samples using the DNeasy PowerSoil Kit (Qiagen, 47014) according to the manufacturer’s protocol, with mechanical disruption in bead-beating tubes. DNA quantity and purity were assessed by Qubit dsDNA BR Assay (Thermo Fisher Scientific, Q32853) and NanoDrop spectrophotometry, with purity ratios at A260/280 of approx. 1.8 and at A260/230 between 1.8-2.2. DNA integrity was confirmed by 1% agarose gel electrophoresis. Full-length 16S rRNA genes were amplified using a degenerate primer set enabling broad bacterial detection, including *Bifidobacterium* spp., and barcoded with Oxford Nanopore Technologies (ONT) native barcodes following the method of Dommann et al.^46^. Amplicons were pooled in equal concentrations and processed for sequencing on a MinION device using the Native Barcoding Kit 96 V14 with Flongle flow cells R10.4.1, following manufacturer protocols. Sequencing runs lasted ∼24 h.

Demultiplexing and basecalling were performed with Dorado (v0.2.4) using the dna_r10.4.1_e8.2_400bps_hac@v4.1.0 model. Reads were quality-filtered and length-trimmed (NanoFilt v2.8.0; 1,300 to 1,800 bp; Q-score >9), with MultiQC (v1.11) used for run quality assessment. Inner PCR barcode demultiplexing and trimming were conducted with seqkit (v2.6.1) and custom scripts based on the 2BC barcode combination. Taxonomic assignment was performed with Emu (v3.4.5) using the default rrnDB v5.6 and NCBI 16S RefSeq (September 2020) database, producing species-level annotations based on full-length 16S matches. The Nextflow pipeline was built using singularity containers from https://depot.galaxypro-ject.org/singularity/.

### Metabolomics

Quantitative profiling of SCFAs in fecal samples was performed by Metabolon. Briefly, samples were spiked with stable labeled internal standards, followed by homogenization and protein precipitation using an organic solvent. Extracted samples were subjected to LC-MS/MS analysis using an Agilent 1290/SCIEX QTRAP 5500 system equipped with a C18 reversed-phase UHPLC column. Data acquisition was carried out in negative mode using electrospray ionization. The peak area of each analyte was measured against the peak area of the corresponding internal standard. Quantification was performed using weighted linear least squares regression analysis generated from the spiked calibration standard prepared concurrently with the study samples. Raw LC-MS/MS data was collected using SCIEX Analyst Software 1.7.3 and processed using SCIEX OS-MQ software v3.1.6.

## Data analysis

All data were analyzed using GraphPad Prism 10 (GraphPad Software, La Jolla, CA, USA), unless otherwise specified. Model assumptions of normality and homogeneous variance were assessed by analysis of the model residuals. Data were analyzed by Student’s unpaired *t* test or Mann-Whitney test when comparing two groups or by one-way analysis of variance (ANOVA) with Tukey’s post-test or Kruskal-Wallis test with Dunn’s post-test when comparing three or more groups. Experimental group was considered statistically significant if the fixed effect *F* test *P* value was ≤0.05. Post hoc pairwise comparisons between experimental groups were made using Tukey’s honestly significant difference multiple-comparison test. A difference between experimental groups was taken to be significant if the *P* value was less than or equal to 0.05 (\**P* < 0.05; \*\**P* < 0.01; \*\*\**P* < 0.001; \*\*\*\**P* < 0.0001).

## Acknowledgments

We thank the animal facility of Swiss TPH for their support with housing Wildings. We would like to acknowledge the support of the ETH genomics platform for help with analysis of our single cell sequencing dataset, in particular Dr Noe Falko. Many thanks to Prof. Daniela Finke for her valuable comments on our manuscript. We acknowledge the use of Novomagic for analysis of our bulk RNA sequencing dataset. We thank the IMCF, Biozentruum facility for providing support for imaging. We thank Prof. David Voehringer for sharing ILC2 staining protocol and Prof. Katharina Röltgen for support with MSD analysis.

## Author contributions

Conceptualization: TB, SRo

Methodology: RD, DO, VT, AB, SRe, AS, CP, NP, JD, PS, SRu, SPR, TB

Investigation: RD, DO, VT, AB, SRe, AS, CP, NP, JD, PS, SRu, SPR, TB

Funding acquisition: TB, SPR

Supervision: TB, SPR, PS, JD

Writing - original draft: TB, DO, RD

Writing - review & editing: TB, DO, SRu, SPR, PS, JD

---

**Fig S1.**
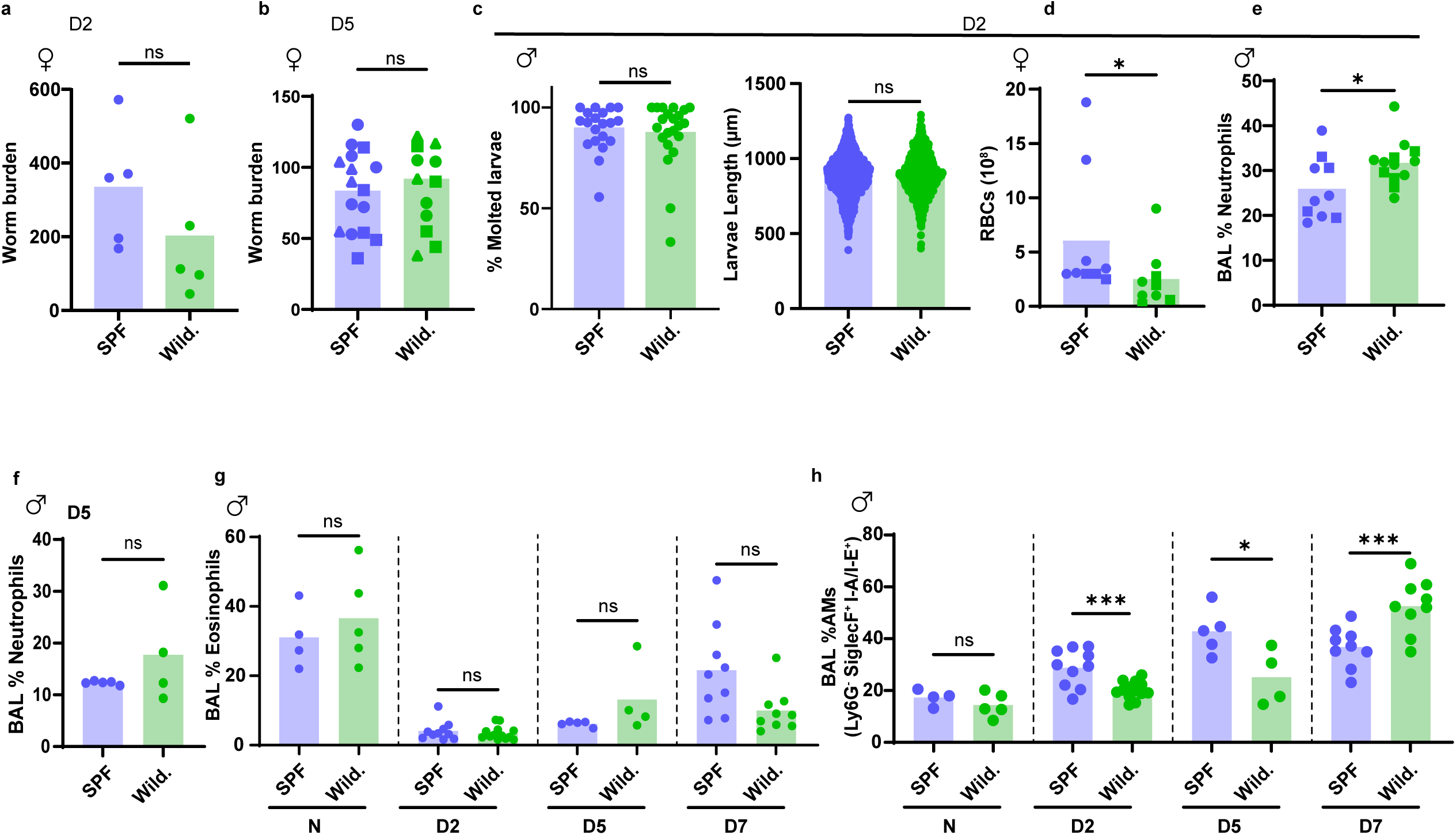
*N. brasiliensis* establishment in the lungs of wildlings mirror SPF response. Wildlings and SPF mice were infected by subcutaneous infection with 500 L3 of *N. brasiliensis,* for 0, 2 or 5 days (Naïve, D2-5). Worm burden was measured by baermann of the (**A**) lungs 2 days post-infection and in the (**B**) small intestine 5 days post-infection.**(C)** Lung larvae harvested by bronchoalveolar lavage and baerman were pooled and measured based on images. Their molting stage was assessed based on morphology. (**D**) 2 days post-infection, bronchoalveolar lavage of the lungs was performed, and red blood cells (RBCs) counted to assess hemorrhaging. **(E-H)** Flow cytometry of BAL samples was performed to quantify the live proportion of immune cell subpopulations in the airways of each mouse. CD45^+^ Ly6G^+^ Gr1^+^ neutrophils were quantified at D2 and D5 post-infection (**E**) and (**F**) respectively. CD45^+^ LyG^−^ SiglecF^+^ I-A/I-E^−^ SCC^high^ eosinophils **(G)** and CD45^+^ LyG^−^ SiglecF^+^ I-A/I-E^+^ alveolar macrophages (AMs) (**H**) were quantified in naive uninfected (N) mice, and at 2, 5, and 7 days post-infection (D2-7). Each dot represents one mouse. D2 time points are pooled from 3 independent experiments (n=4-5); D5 and Naive are issued from 1 experiment (n=4-5). When possible, different symbols are used between biological repeats.

**Fig S2.**
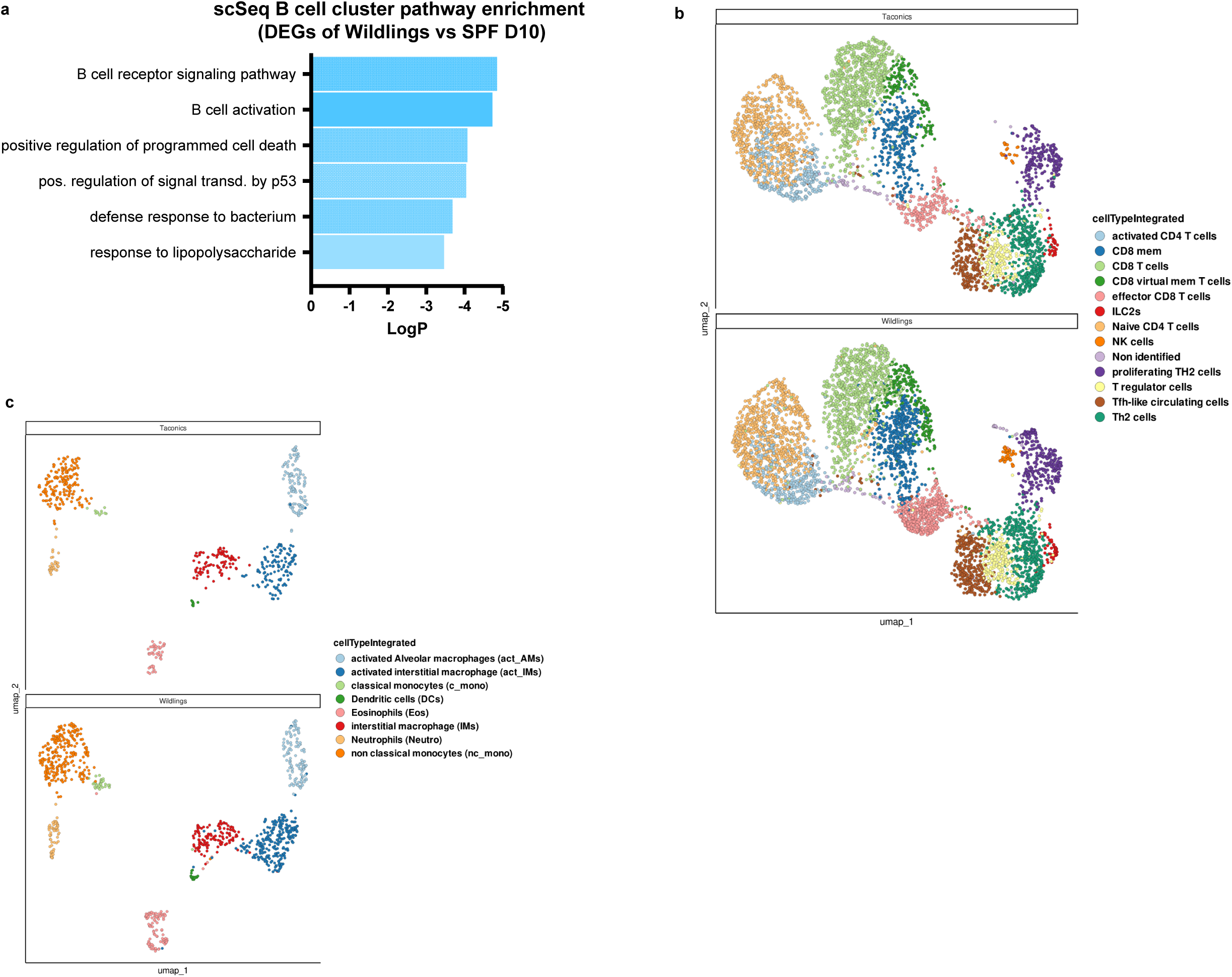
Single cell sequencing of the lung immune response to *N. brasiliensis* in wildlings and SPF mice. Wildlings and SPF female mice were infected by subcutaneous infection with 500 L3 of *N. brasiliensis,* 10 days. Lungs cells were isolated by digestion from naïve or 10 days infected mice and single cell RNA sequencing was performed on a BD Rhapsody (n=4, per mouse type). **(A)** Pathway enrichment analysis performed in Metascape for differentially expressed genes between SPF and wildlings in the overall B cell cluster. **(B-C)** UMAP visualization of all cells clustered by Seurat in wildlings and SPF mice. Cells were sorted in “lymphoid” **(B)** and “myeloid” **(C)** for further analysis.

**Fig S3.**
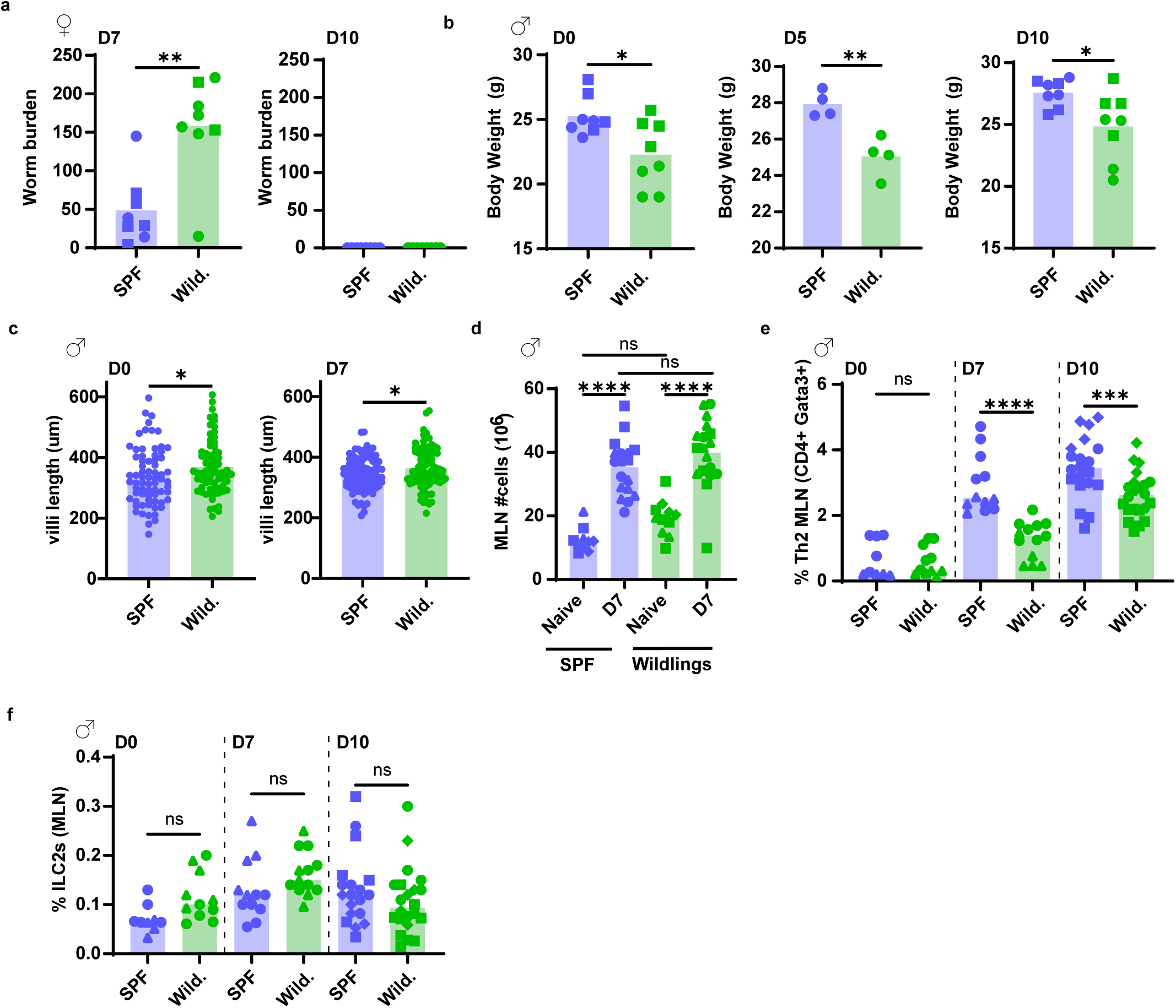
Wildlings have a delayed intestinal expulsion of *N. brasiliensis*. Wildlings and SPF mice were infected by subcutaneous infection with 500 L3 of *N. brasiliensis, 7-*10 days. Worm burden was measured by baermann of the (**A**) small intestine 7 or 10 days post-infection. Data pooled from 3 independent experiments (n= 4-8) **(B)** mice body weight were measured in several mouse cohorts in naive or infected mice (D5, D10). Data pooled from 2 independent experiments (n=4-5). **(C)** villi length was measured on swiss roll in the second centimeter of duodenum in naive or D7 infected mice. Data representative of 2 independent experiments, each data point represents a villus, with 10 villi counted per mouse (n=4-5). **(D)** Mesenteric Lymph Nodes from a complete chain (MLN) were crushed and total cells were enumerated. **(E-F)** Flow cytometric analysis of Th2 cells (K, CD45^+^ CD4^+^ Gata-3^+^) and ILC2s (L, CD45^+^ Lin^−^ KLRG1^+^ Gata-3^+^) in the MLN of naïve mice or mice in early or late phase of expulsion. 1 experiment, representative of 2 more (n=4-5). When possible, different symbols are used between biological repeats.

**Fig S4.**
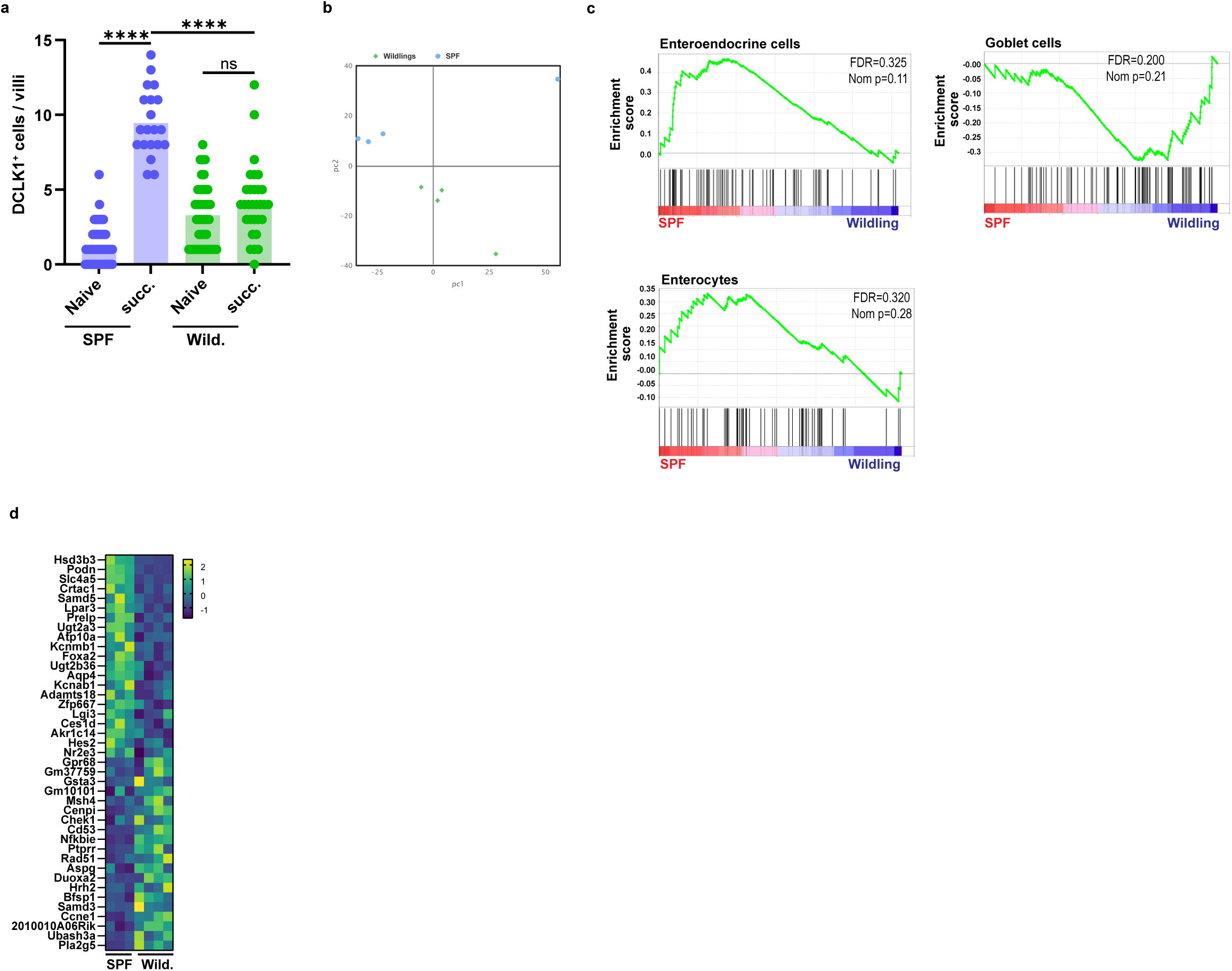
Characterization of the constrained intestinal epithelial cell response in wildlings. **(A)** Wildlings and SPF mice were supplemented with succinate in the drinking water for 7 days. Tuft cells were quantified by immunofluorescence analysis of DCLK1 in the small intestine villi. **(B-C)** Bulk RNA sequencing the first centimeter of duodenum in wildlings or SPF mice 7 days post-infection with 500 L3 of *N. brasiliensis* (n=4). Principal component analysis **(B)**. Enrichment plots generated by GSEA analysis of tuft cells as defined in Habes et al **(C). (D)** Heatmap of tuft-p genes involved in the GSEA analysis described in O. The first 20 genes were considered leading genes. When possible, different symbols are used between biological repeats.

**Fig S5.**
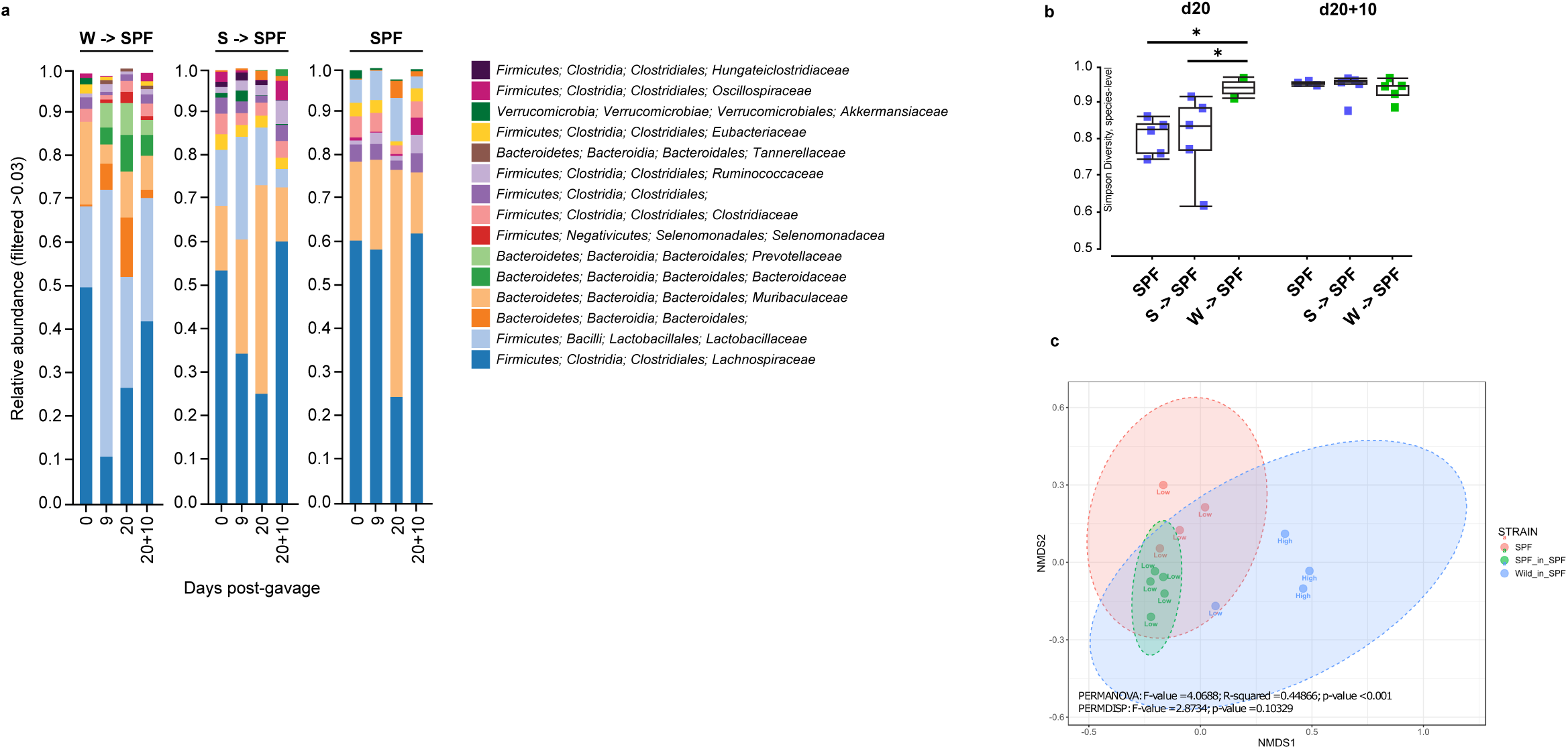
Bacterial microbiome characterization after caecal microbiota transfer at adult age. Caecal material from Wildlings or SPF was transplanted by oral gavage into SPF mice. One SPF group was left untreated. After 20 days post colonization, all mice were infected with 500 L3 of *N. brasiliensis* for 7-9 days. Fecal pellets from all mice were collected over 27-29 days and long read16S rRNA gene profiling by nanopore was performed. **(A)** Relative abundance of the bacterial Family of all mice, before and during colonization, and post-infection. **(B)** a-diversity calculated with a Simpson index, at the species level. **(C)** PCA based on relative abundance of 20 days post-colonization and 7 days post-infection with Wildlings or SPF material. Data was segregated by egg burden: Low = 0-1 eggs at day 30; High = 1+ (9-24) eggs at day 30

